# A Wnt-responsive fibrocartilage progenitor system coordinates postnatal mandibular condylar cartilage growth

**DOI:** 10.64898/2026.03.25.714159

**Authors:** Toshihiro Inubushi, Renshiro Kani, Yutaro Tanida, Yu Usami, Tomoaki Iwayama, Wu Deyang, Jun-Ichi Sasaki, Jingyu Ye, Shinnosuke Kusano, Yuki Shiraishi, Hiroshi Kurosaka, Dragana Kopanja, Masahide Takedachi, Shinya Murakami, Takashi Yamashiro

**Affiliations:** Department of Orthodontics and Dentofacial Orthopedics, The University of Osaka Graduate School of Dentistry, Osaka, Japan; Department of Oral and Maxillofacial Pathology, The University of Osaka Graduate School of Dentistry, Suita, Japan; Department of Periodontology and Regenerative Dentistry, The University of Osaka, Graduate School of Dentistry.; Department of Dental Biomaterials, The University of Osaka Graduate School of Dentistry, Osaka, Japan; Department of Biochemistry and Molecular Genetics, University of Illinois College of Medicine

**Author notes:** Corresponding author: Toshihiro Inubushi, Department of Orthodontics and Dentofacial Orthopedics, The University of Osaka Graduate School of Dentistry. 1-8 Yamada-oka, Suita, Osaka, 565-0871, Japan. These authors contributed equally to this work (Toshihiro Inubushi, Renshiro Kani).

**Keywords:** Wnt/β-catenin, Temporomandibular joint, fibrocartilage, Foxm1

## Abstract

Postnatal growth of the mandibular condyle requires coordinated expansion of fibrocartilage and production of chondrocytes, yet the cellular populations that organize this process remain incompletely defined. Here we identify a Wnt-responsive fibrocartilage progenitor population that contributes to postnatal mandibular condylar cartilage growth. Using a direct Wnt activity reporter (*R26-WntVis*), inducible genetic lineage tracing (*Axin2^CreERT2^*), and single-cell transcriptomics, we define a Wnt-enriched progenitor-like cluster localized predominantly within the fibrocartilage zone. Lineage tracing demonstrates that Axin2-lineage cells expand laterally within fibrocartilage and generate vertically aligned chondrocytes in the chondrocartilage compartment, indicating bidirectional growth contribution *in vivo*.

Conditional ablation of β-catenin in Axin2-lineage cells results in depletion of the fibrocartilage compartment and premature activation of chondrogenic differentiation programs, whereas constitutive β-catenin activation disrupts compartmental organization without enhancing proliferation. Mechanistically, we identify Foxm1 as a Wnt-associated proliferative mediator enriched in fibrocartilage, and genetic reduction of Foxm1 cooperates with β-catenin deficiency to impair condylar growth. In parallel, β-catenin loss derepresses TGF-β–Smad signaling and enhances chondrogenic differentiation, indicating that canonical Wnt activity coordinates proliferative maintenance while restraining lineage commitment within the same cellular compartment.

Together, these findings identify a Wnt-responsive fibrocartilage progenitor system that regulates postnatal mandibular condylar cartilage growth by coupling Foxm1-associated proliferative maintenance with suppression of TGF-β–dependent chondrogenic differentiation during temporomandibular joint development.

**Graphical abstract:** Wnt-responsive fibrocartilage progenitors coordinate postnatal mandibular condylar cartilage growth through Foxm1-dependent proliferative maintenance and suppression of TGF-β–driven chondrogenic differentiation.

## Introduction

Postnatal skeletal growth requires coordinated regulation of progenitor expansion and differentiation across distinct anatomical growth centers to ensure proper tissue morphogenesis and function (Enlow & Hans, 1996; McNamara, 1980). Among these, the mandibular condylar cartilage (MCC) serves as a principal driver of mandibular elongation and temporomandibular joint (TMJ) morphogenesis. Disruption of MCC growth is closely associated with condylar hypoplasia, mandibular retrusion or overgrowth, and degenerative TMJ disease (Arnett, Milam, & Gottesman, 1996; Bryndahl, Eriksson, Legrell, & Isberg, 2006; Schellhas, Pollei, & Wilkes, 1993; Tanaka, Detamore, & Mercuri, 2008). Experimental models further demonstrate that dysregulated signaling within the condylar cartilage results in impaired fibrocartilage maintenance and progressive joint degeneration (Embree et al., 2016). Despite its clinical relevance, the cellular and molecular mechanisms that sustain postnatal MCC growth remain incompletely understood.

Histologically, MCC is organized into a superficial zone, an underlying fibrocartilage compartment enriched in collagen type I, and a deeper chondrocartilage region characterized by collagen type II–producing chondrocytes (Detamore & Athanasiou, 2003; Shibukawa et al., 2007). Unlike the hyaline cartilage of long bone growth plates, MCC represents a form of secondary cartilage within a fibrocartilage-dominant environment that supports both lateral tissue expansion and vertical chondrocyte production. This unique architecture suggests that proliferative maintenance and differentiation must be tightly coordinated within the fibrocartilage compartment. In long bones, skeletal stem and progenitor cells residing in the resting zone of the growth plate have been well characterized (Chan et al., 2015; Mizuhashi et al., 2018; Park et al., 2012). These populations contribute to cartilage, bone, and stromal lineages during growth and regeneration and are defined by distinct transcriptional programs and long-term lineage tracing approaches. In contrast, equivalent progenitor populations within MCC remain less clearly defined. Several studies have proposed progenitor-like cells within the fibrocartilage compartment based on mesenchymal markers such as Col1 or α-smooth muscle actin (Acri et al., 2019; Kalajzic et al., 2008; Robinson, O’Brien, Chen, & Wadhwa, 2015). However, these markers label heterogeneous populations and do not resolve lineage hierarchy or long-term functional contribution, leaving the identity and regulatory mechanisms of MCC progenitors unresolved.

Canonical Wnt/β-catenin signaling is a central regulator of progenitor proliferation and lineage specification across multiple tissues (Bhavanasi & Klein, 2016; Holland, Klaus, Garratt, & Birchmeier, 2013; Nusse & Clevers, 2017). In skeletal mesenchymal cells, Wnt signaling promotes osteogenic commitment while restraining chondrogenic differentiation (Day, Guo, Garrett-Beal, & Yang, 2005; Hill, Später, Taketo, Birchmeier, & Hartmann, 2005) and regulates cell cycle progression through activation of Cyclin D1 and repression of cell cycle inhibitors (Gartel et al., 2001; Tetsu & McCormick, 1999). Wnt signaling also interfaces with metabolic growth pathways through GSK3-dependent mechanisms (Inoki et al., 2006). In several contexts, canonical Wnt activity functionally antagonizes TGF-β–Smad signaling to influence lineage commitment and differentiation programs (Guo & Wang, 2009). However, how Wnt signaling operates within the structurally distinct and fibrocartilage-dominant environment of MCC remains unclear, particularly with respect to how proliferative expansion and differentiation are coordinated within the same cellular compartment.

Recent studies have begun to explore TMJ progenitor populations. *Lgr5*-positive cells contribute to synovial and disc components of the joint but do not directly generate condylar chondrocytes (Ruscitto et al., 2023). More recently, *FSP1/S100A4*-positive fibrocartilage cells were implicated in TMJ integrity and proposed to reside within a Wnt-suppressive niche (Tuwatnawanit et al., 2025). However, these conclusions were based largely on static marker expression and constitutive genetic models without direct assessment of dynamic Wnt transcriptional activity or long-term lineage contribution within MCC. Thus, whether MCC contains a defined Wnt-responsive proliferative population that coordinates fibrocartilage expansion and chondrocartilage differentiation remains an open question.

Here, we sought to determine whether MCC harbors a functionally defined Wnt-responsive proliferative population and to define its contribution to postnatal condylar growth. Using a direct Wnt activity reporter (*R26-WntVis*), inducible genetic lineage tracing (*Axin2^CreERT2^*), single-cell transcriptomics, and genetic perturbation approaches, we identify a Wnt-responsive fibrocartilage-resident population that contributes to both lateral fibrocartilage expansion and vertical chondrocartilage formation. Functional analyses demonstrate that physiological β-catenin activity is required to sustain proliferative capacity while limiting premature TGF-β–dependent chondrogenic differentiation. We further implicate Foxm1 as an important mediator associated with Wnt-dependent proliferative programs in MCC, consistent with prior evidence of cooperative interactions between β-catenin and Foxm1 in other systems (Zhang et al., 2011). Together, our findings provide a mechanistic framework for understanding how proliferative maintenance and differentiation progression are coordinated within the fibrocartilage compartment during postnatal mandibular condyle growth.

## Results

### Identification of Wnt-responsive cells in postnatal mandibular condylar cartilage

To identify Wnt-responsive cell populations that may regulate postnatal mandibular condylar cartilage growth, we first examined the spatial distribution of canonical Wnt/β-catenin signaling using the Wnt reporter mouse line *R26-WntVis*, in which H2B–EGFP expression is driven by seven tandem TCF/LEF binding elements (Fig. 1A). At postnatal day 14 (P14), sagittal sections revealed prominent H2B-EGFP–positive nuclei within the superficial and fibrocartilage zones, whereas reporter activity was largely absent from the deeper chondrocartilage compartment (Fig. 1B). These observations indicate that Wnt reporter activity is enriched in the superficial and fibrocartilage compartments of the postnatal mandibular condyle.

**Figure 1.**
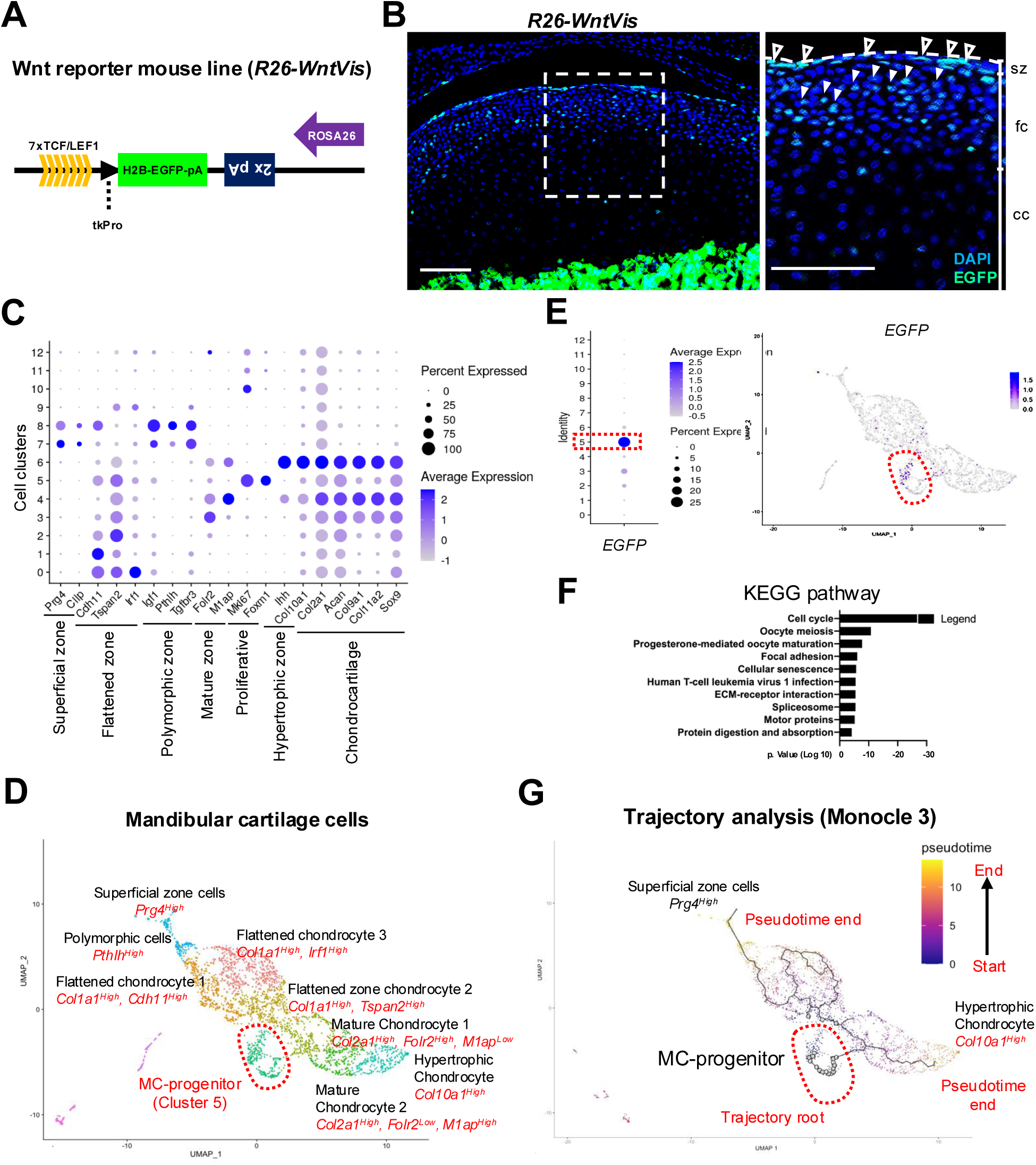
Identification of a Wnt-responsive progenitor population in postnatal mandibular condylar cartilage. (A) Schematic representation of the *R26-WntVis* reporter. H2B–EGFP expression is driven by seven tandem TCF/LEF binding elements upstream of a minimal thymidine kinase promoter, enabling visualization of canonical Wnt/β-catenin signaling activity. (B) Sagittal frozen sections from *R26-WntVis* mice at postnatal day 14 (P14) were analyzed by fluorescence microscopy. H2B–EGFP–positive nuclei are enriched in the superficial (sz; open arrowheads) and fibrocartilage (fc; closed arrowheads) zones and largely absent from the deeper chondrocartilage (cc) zone, indicating spatial restriction of canonical Wnt activity to the upper compartments. Dashed boxes indicate regions shown at higher magnification. Scale bar, 100 µm. Data are representative of three biologically independent mice. (C) Single-cell RNA sequencing was performed on enzymatically dissociated mandibular condylar cartilage from P16 *R26-WntVis* mice. Uniform Manifold Approximation and Projection (UMAP) visualization identified nine transcriptionally distinct clusters following quality control filtering and shared nearest neighbor clustering. (D) Feature plots show expression of representative marker genes used to annotate fibrocartilage (*Col1a1*) and chondrocartilage (*Col2a1*) populations, enabling classification of major cartilage compartments. (E) Feature plot and dot plot analyses were used to examine H2B–EGFP reporter expression across clusters. Cluster 5 shows marked enrichment of H2B–EGFP–positive cells and was designated as the MC-progenitor cluster, indicating a Wnt-responsive progenitor-like population. (F) KEGG pathway enrichment analysis was performed on differentially expressed genes in the MC-progenitor cluster relative to differentiated cartilage clusters. Cell-cycle–associated pathways are significantly enriched, consistent with a proliferative transcriptional state. (G) Trajectory inference was performed using Monocle3 to assess lineage relationships among clusters. The MC-progenitor cluster (cluster 5) is positioned at the root of a bifurcating trajectory leading toward fibrocartilage and chondrocartilage lineages, supporting its role as an upstream progenitor population.

### Single-cell transcriptomic identification of an MC-progenitor cluster

P16 was selected for transcriptomic analysis because this stage corresponds to active postnatal growth of the mandibular condyle, characterized by expansion of the fibrocartilage compartment. To molecularly characterize Wnt-responsive populations, we performed single-cell RNA sequencing (scRNA-seq) on enzymatically dissociated mandibular condylar cartilage pooled from two independent P16 *R26-WntVis* mice. After quality control filtering, uniform manifold approximation and projection (UMAP) analysis resolved multiple transcriptionally distinct clusters, including nine cartilage-associated clusters (clusters 0–8) and several minor clusters corresponding to non-cartilage cell populations (Fig. 1C). These clusters were annotated based on established marker gene expression profiles (Extended Data Fig. 2). Cells segregated into major fibrocartilage (*Col1a1⁺*) and chondrocartilage (*Col2a1⁺*) populations, together with smaller clusters representing endothelial and immune cells (Fig. 1C, D). Notably, one discrete cluster (Cluster 5) displayed reduced expression of differentiated cartilage markers while exhibiting strong enrichment of *H2B–EGFP* transcripts (Fig. 1E). Based on these features, we designated this population as the MC-progenitor cluster, as these cells exhibited (i) reduced expression of lineage-committed cartilage markers, (ii) enrichment of canonical Wnt reporter transcripts, and (iii) a transcriptional profile consistent with a progenitor-like state relative to adjacent fibrocartilage and chondrocartilage populations. Gene ontology and KEGG pathway analyses revealed enrichment of cell-cycle–associated pathways in the MC-progenitor cluster, consistent with a proliferative transcriptional program (Fig. 1F). Quality control filtering removed cells with low transcript counts, high mitochondrial transcript content, or abnormally high gene numbers, yielding a high-quality dataset for downstream analyses (Extended Data Fig. 1).

### Trajectory and RNA velocity analyses position the MC-progenitor cluster at the apex of a bifurcating hierarchy

To investigate lineage relationships among mandibular condylar cartilage cell populations, trajectory analysis was performed using Monocle3. Pseudotime ordering positioned the MC-progenitor cluster (Cluster 5) at the origin of an inferred bifurcating trajectory extending toward both fibrocartilage and chondrocartilage lineages (Fig. 1G). Based on its placement at the earliest pseudotime position, Cluster 5 was defined as the trajectory root. To independently assess lineage directionality, RNA velocity analysis was performed using scVelo. Projection of velocity vectors onto the UMAP embedding revealed coherent directional flows emanating from the MC-progenitor cluster toward both fibrocartilage and chondrocartilage populations, consistent with the differentiation trajectories inferred by Monocle3 (Extended Data Fig. 3). Together, these transcriptional trajectory and RNA velocity analyses support a model in which the MC-progenitor cluster occupies the apex of a bifurcating lineage hierarchy within postnatal mandibular condylar cartilage. These findings further suggest that Wnt-responsive progenitor populations within the fibrocartilage compartment may contribute to both fibrocartilage maintenance and chondrocartilage production during postnatal growth, a possibility we next tested using genetic lineage tracing *in vivo*.

### Axin2-lineage tracing demonstrates dual-lineage contribution of Wnt-responsive cells *in vivo*

To test whether the Wnt-responsive progenitor population identified by single-cell transcriptomic analyses contributes to postnatal mandibular condylar cartilage growth in vivo, we performed inducible genetic lineage tracing using *Axin2^CreERT2^*mice. Tamoxifen was administered at postnatal day 14 (P14), and lineage-labeled cells were analyzed at defined time points after induction. At 2 days post-induction (P16), ZsGreen-positive (ZsG⁺) cells were predominantly localized to the superficial and upper fibrocartilage zones (Fig. 2A). At 7 days post-induction (P21), ZsG⁺ cells remained enriched within the proliferative fibrocartilage compartment and formed small expanding clusters. By 28 days post-induction (P42), ZsG⁺ cells extended vertically into the chondrocartilage zone, forming columnar arrays that included hypertrophic chondrocytes, while also expanding laterally within the fibrocartilage layer. These findings demonstrate that Axin2-lineage cells contribute to both fibrocartilage and chondrocartilage compartments during postnatal growth. To assess the maximal contribution of Axin2-lineage cells, mice received three consecutive tamoxifen injections at P14 and were analyzed at 28 days post-induction (P42). Under these conditions, ZsGreen labeling encompassed nearly all fibrocartilage cells and a large proportion of mature chondrocytes, indicating that endogenous Wnt-responsive cells represent a major cellular source for postnatal mandibular condylar cartilage expansion.

**Figure 2.**
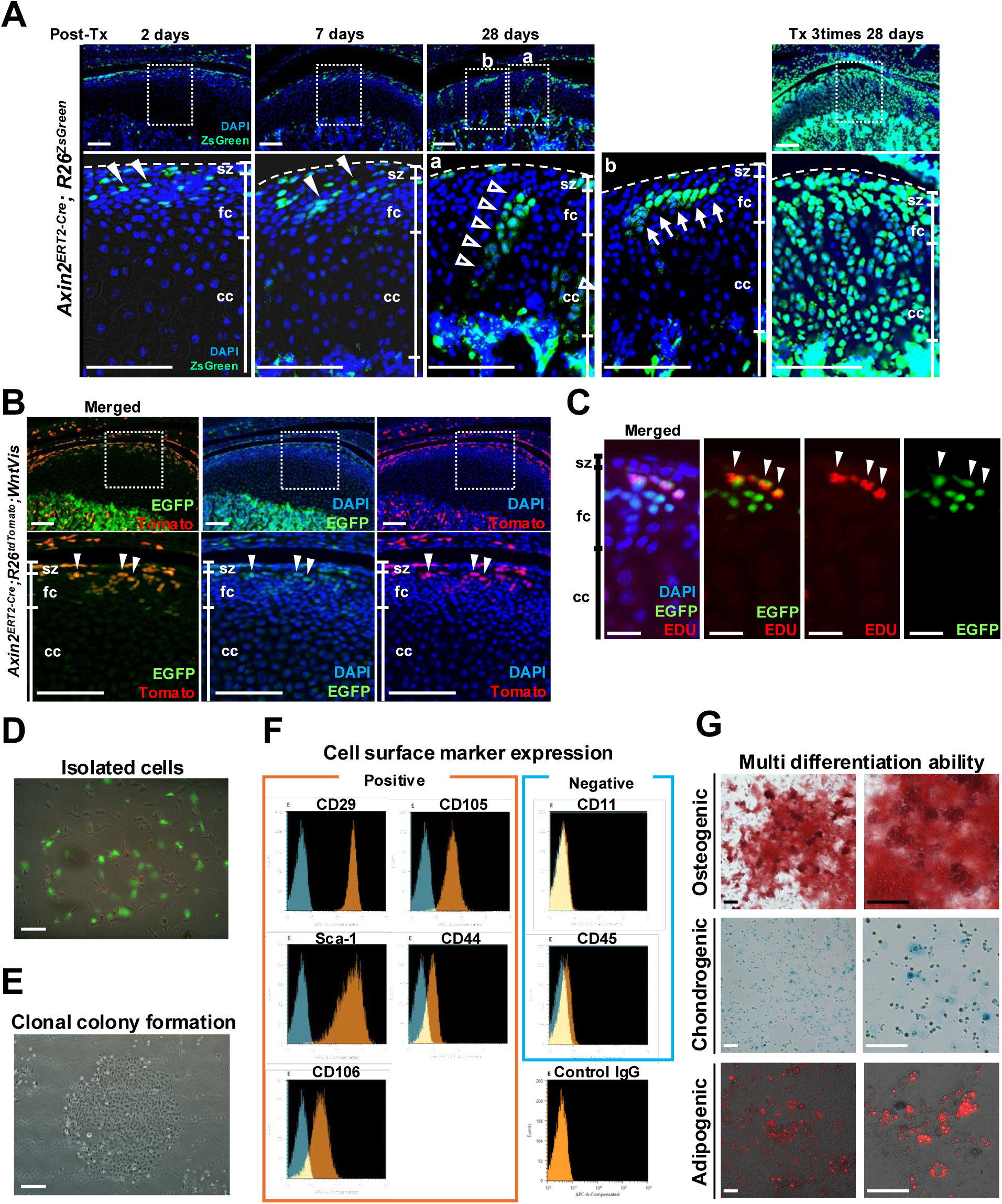
Lineage tracing reveals bidirectional contributions of Wnt-responsive cells to condylar cartilage growth. (A) Inducible lineage tracing was performed using *Axin2^CreERT2^;Rosa26^ZsGreen^*mice. Tamoxifen was administered at postnatal day 14 (P14), and ZsGreen-positive (ZsG⁺) cells were analyzed at defined time points after induction (P16, P21, and P42). At 2 days post-induction (P16), ZsG⁺ cells are predominantly localized within the superficial and fibrocartilage zones. At 7 days post-induction (P21), ZsG⁺ cells expand within the fibrocartilage compartment and form small clusters. By 28 days post-induction (P42), ZsG⁺ cells extend vertically into the chondrocartilage zone and form columnar arrays (open arrowheads), while cells expanding laterally within the fibrocartilage layer are indicated by arrows. Following repeated tamoxifen administration, ZsG⁺ cells populate most of the fibrocartilage layer and a large proportion of chondrocytes, indicating both lateral and vertical contributions during postnatal growth. The lower panels show higher-magnification views of the regions outlined by dashed boxes in the upper panels. Closed arrowheads indicate H2B–EGFP–positive cells within the fibrocartilage (fc) zone. Representative images from n = 3 biologically independent mice per time point are shown. Scale bar, 100 μm. (B) To compare canonical Wnt signaling activity with Axin2-lineage labeling *in vivo*, *Axin2^CreERT2^;Rosa26^Tomato^;R26-WntVis* mice were analyzed two days after tamoxifen administration. H2B–EGFP–positive nuclei (green), indicating active canonical Wnt signaling, show substantial spatial overlap with Tomato-labeled Axin2-lineage cells (red) within the superficial (sz) and fibrocartilage (fc) zones. Closed arrowheads indicate H2B–EGFP–positive cells within the fibrocartilage zone, which co-express Tomato, demonstrating concordance between Wnt activity and lineage labeling. Nuclei are counterstained with DAPI (blue). Representative images from three biologically independent mice are shown. Scale bar, 100 μm. (C) To assess proliferative dynamics, EdU label-retention analysis was performed. EdU was administered 28 days prior to tissue harvest, and EdU-retaining cells were detected within the upper fibrocartilage layer. Closed arrowheads indicate EdU-retaining cells that co-express H2B–EGFP, identifying a slow-cycling Wnt-responsive subpopulation. Scale bar, 20 μm. (D) H2B–EGFP–positive cells were isolated from mandibular condylar cartilage of P14 R26-WntVis mice by fluorescence-activated cell sorting (FACS). (E) Clonal colony formation assays were performed following *in vitro* culture. Isolated cells form colonies, indicating clonogenic potential. (F) Flow cytometry analysis was performed to characterize cell surface marker expression. Isolated cells express mesenchymal stromal cell–associated markers (CD29, CD105, Sca-1, CD44, CD106) and lack hematopoietic markers (CD45 and CD11), indicating a mesenchymal stromal phenotype. (G) Isolated Wnt-responsive cells were cultured under osteogenic, chondrogenic, or adipogenic differentiation conditions. Cells exhibit multilineage differentiation capacity, as evidenced by positive staining with Alizarin Red (osteogenesis), Alcian Blue (chondrogenesis), and Oil Red O (adipogenesis), respectively. Scale bar, 100 μm. Abbreviations: sz, superficial zone; fc, fibrocartilage zone; cc, chondrocartilage zone.

### *In vivo* correspondence between Wnt reporter activity and Axin2-lineage labeling

To directly compare ongoing canonical Wnt signaling activity with Axin2-lineage labeling *in vivo*, we generated *Axin2^CreERT2^;Rosa26^Tomato^;R26-WntVis* mice and performed double fluorescence analysis following tamoxifen administration. Two days after tamoxifen induction, substantial spatial overlap was observed between H2B-EGFP–positive nuclei, indicating active canonical Wnt signaling, and Tomato-labeled Axin2-lineage cells within the superficial and fibrocartilage zones of the mandibular condyle (Fig. 2B). Although the two signals showed strong spatial correspondence, complete concordance was not expected because the reporters capture distinct biological readouts. The WntVis reporter reflects real-time TCF/LEF transcriptional activity, whereas Axin2-driven Cre recombination permanently labels cells that have experienced endogenous Wnt signaling following tamoxifen-induced recombination. Despite these mechanistic differences, the substantial spatial overlap indicates that Axin2-lineage tracing largely captures the endogenous Wnt-responsive compartment corresponding to the MC-progenitor population identified by single-cell transcriptomics. These findings further support that the Axin2-lineage population corresponds to the endogenous Wnt-responsive progenitor compartment identified by the Wnt reporter and single-cell transcriptomic analyses.

### EdU label retention identifies a slow-cycling Wnt-responsive subpopulation

To examine proliferative dynamics *in vivo*, we performed an EdU label-retention assay. Twenty-eight days after EdU administration, a small population of EdU-retaining cells was detected within the upper fibrocartilage layer of the mandibular condyle. A subset of these cells co-expressed H2B–EGFP in *R26-WntVis* mice (Fig. 2C), indicating that a fraction of Wnt-responsive cells exhibits slow-cycling behavior. These findings are consistent with the presence of a slow-cycling Wnt-responsive subpopulation that may contribute to long-term maintenance of the fibrocartilage compartment.

### Functional characterization of isolated Wnt-responsive cells

To further assess the functional properties of Wnt-responsive cells, H2B–EGFP–positive cells were isolated by fluorescence-activated cell sorting (FACS) from mandibular condylar cartilage at P14 (Fig. 2D). Expanded cells expressed mesenchymal stromal cell–associated markers (CD29, CD105, Sca-1, CD44, and CD106) and lacked hematopoietic markers (CD45 and CD11) (Fig. 2F). Isolated cells formed clonal colonies *in vitro* (Fig. 2E) and, under defined culture conditions, exhibited osteogenic, chondrogenic, and adipogenic differentiation capacity (Fig. 2G), consistent with multilineage mesenchymal differentiation potential *in vitro*. Collectively, these complementary single-cell, lineage-tracing, and functional analyses identify a Wnt-responsive fibrocartilage progenitor population that contributes to both fibrocartilage maintenance and chondrocartilage production during postnatal mandibular condylar cartilage growth.

### β-catenin signaling maintains proliferative fibrocartilage and restrains premature chondrogenic differentiation

To determine the functional requirement of canonical Wnt/β-catenin signaling in Axin2-lineage cells, we generated *Axin2^CreERT2^;Ctnnb1^fl/fl^;Rosa26^ZsGreen^* mice and induced recombination at P14. In this model, Cre-mediated recombination results in irreversible ZsGreen expression from the Rosa26 locus, thereby marking recombined Axin2-lineage cells. Recombination efficiency averaged 23.8 ± 5.0% of fibrocartilage cells (n = 5 biologically independent mice), indicating mosaic recombination within the Axin2-lineage compartment. Histological analysis at P42 revealed a marked reduction of the fibrocartilage compartment compared to controls, whereas the overall chondrocartilage structure remained partially preserved (Fig. 3A). Quantitative analysis confirmed that the fibrocartilage layer thickness was significantly reduced in β-catenin–deficient condyles, whereas the mature chondrocartilage layer thickness showed no significant change (Fig. 3C). Ki67 immunostaining at P21 demonstrated a significant reduction in proliferative cells within ZsGreen-positive (β-catenin–deficient) cells as well as in adjacent ZsGreen-negative cells within the same fibrocartilage region (Fig. 3B, C), which was confirmed by quantitative analysis of Ki67-positive cells. ZsGreen-negative cells were defined as cells lacking detectable reporter expression and therefore served as neighboring non-recombined cells. The reduction of proliferation in these adjacent cells is therefore consistent with a potential non–cell-autonomous effect of β-catenin loss on the surrounding tissue environment. In addition to reduced proliferation, β-catenin–deficient ZsGreen-positive cells extended into the chondrocartilage zone and formed columnar arrangements. As shown in subsequent analyses (Fig. 5), these cells exhibit increased TGF-β–Smad signaling and upregulation of chondrogenic markers, suggesting that loss of β-catenin may promote premature chondrogenic differentiation. Consistent with Ki67 immunostaining, EdU incorporation assays also revealed a significant reduction in proliferating cells in β-catenin–deficient condyles compared with controls, confirming impaired proliferative activity within the fibrocartilage compartment. In contrast, apoptosis was rarely detected in either genotype and showed no significant difference between control and mutant condyles, indicating that the reduction in fibrocartilage thickness primarily reflects impaired cell proliferation rather than increased cell death. Micro–computed tomography (micro-CT) analysis further demonstrated reduced condylar length in β-catenin–deficient mice compared with controls, consistent with the reduction of the fibrocartilage compartment observed histologically (Extended Data Fig. 4). These results collectively suggest that canonical Wnt/β-catenin signaling functions as a regulatory axis that maintains the proliferative fibrocartilage progenitor pool while preventing premature chondrogenic differentiation during postnatal mandibular condylar cartilage growth.

**Figure 3.**
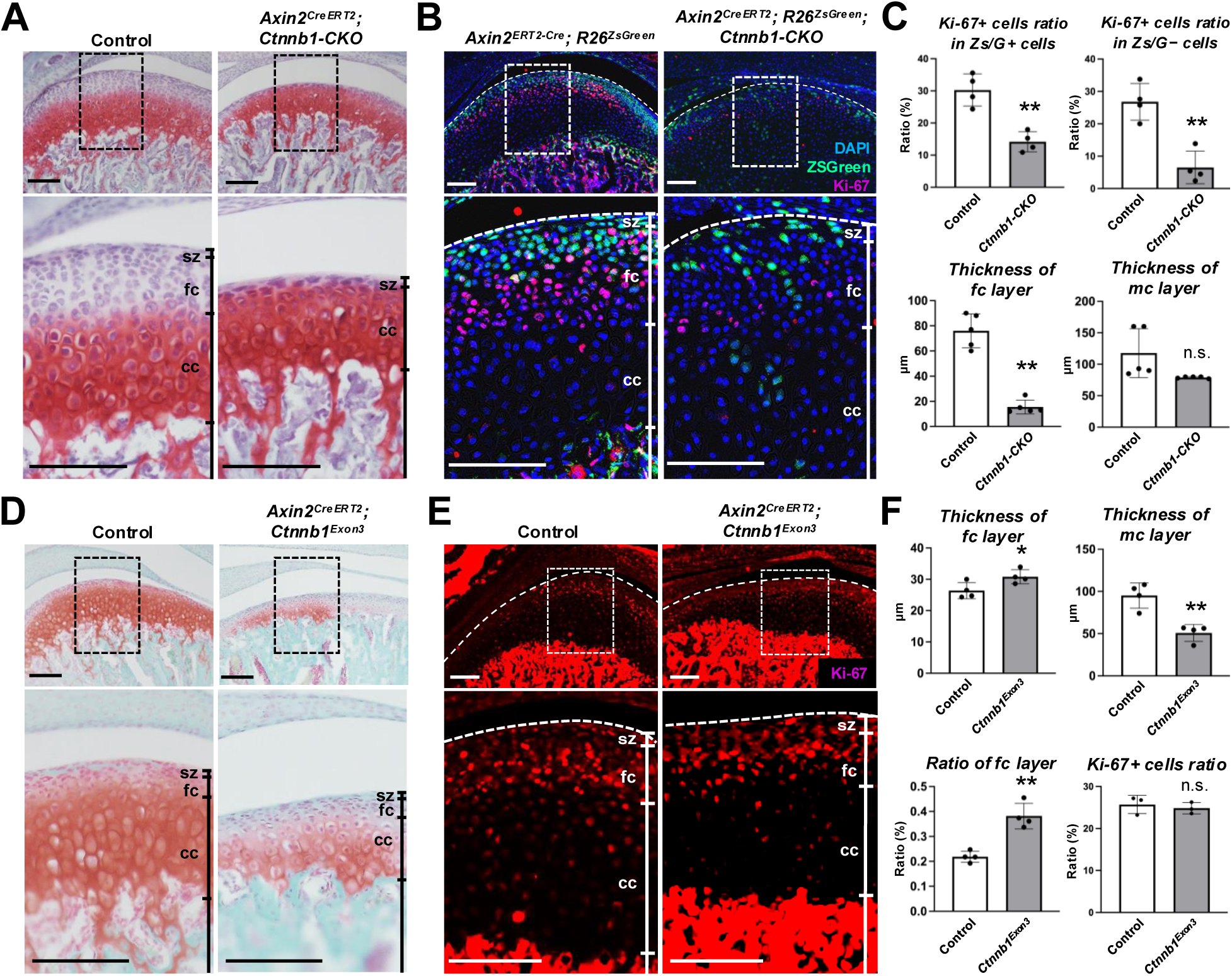
β-catenin maintains fibrocartilage proliferation and prevents premature chondrogenic differentiation. (A) Safranin-O/Fast Green staining was performed on sagittal sections of mandibular condylar cartilage from control and *Axin2^CreERT2^;Ctnnb1^fl/fl^*mice at P42 (28 days after tamoxifen administration). Mutant condyles show marked depletion of the fibrocartilage compartment, whereas the chondrocartilage zone is relatively preserved, indicating impaired maintenance of the fibrocartilage layer upon β-catenin loss. Lower panels show higher magnification views of boxed regions. Scale bar, 100 μm. (B) Immunofluorescence staining for Ki67 was performed in *Axin2^CreERT2^;Ctnnb1^fl/fl^;Rosa26^ZsGreen^* mice and littermate controls at P21 (7 days post-tamoxifen). ZsGreen marks Axin2-lineage cells, enabling comparison between recombined (ZsGreen-positive) and adjacent non-recombined cells within the same tissue. β-catenin–deficient lineage cells exhibit reduced proliferative activity compared with neighboring cells, indicating a cell-autonomous requirement for β-catenin in fibrocartilage proliferation. Scale bar, 100 μm. (C) Quantification of fibrocartilage thickness, chondrocartilage thickness, and Ki67-positive cells in ZsGreen-positive and ZsGreen-negative populations was performed. β-catenin deficiency significantly reduces fibrocartilage thickness and proliferation within Axin2-lineage cells, without markedly affecting the chondrocartilage compartment. Data represent mean ± SD from biologically independent mice (n = 4–5 per genotype). (D) Safranin-O staining was performed on *Axin2^CreERT2^;Ctnnb1^exon3-flox^*mice and littermate controls at P42. Constitutive β-catenin activation results in altered cartilage organization, characterized by expansion of the fibrocartilage domain and reduction of the safranin-O–positive chondrocartilage zone, indicating disrupted compartmental balance. Lower panels show higher magnification views of boxed regions. Scale bar, 100 μm. (E) Ki67 immunostaining was performed in *Axin2^CreERT2^;Ctnnb1^exon3-flox^*mice at P21. Despite expansion of the fibrocartilage compartment, proliferative activity is not increased compared with controls, indicating that β-catenin activation alters tissue organization without proportionally enhancing proliferation. Scale bar, 100 μm. (F) Quantification of fibrocartilage thickness, chondrocartilage thickness, the ratio of fibrocartilage to total cartilage thickness, and Ki67-positive cells in Axin2-lineage cells was performed. β-catenin activation increases fibrocartilage proportion without significantly increasing proliferation. Data represent mean ± SD from biologically independent mice (n = 5 per genotype). Statistical significance was assessed using two-tailed Student’s t-test. n.s., not significant; *P < 0.05; **P < 0.01. Abbreviations: sz, superficial zone; fc, fibrocartilage zone; cc, chondrocartilage zone.

### Constitutive activation disrupts compartmental balance without enhancing proliferation

Conversely, constitutive activation of β-catenin (*Axin2^CreERT2^;Ctnnb1^exon3-flox^*) altered compartmental organization, characterized by expansion of the fibrocartilage zone and reduction of the chondrocartilage domain (Fig. 3D). Quantitative measurements confirmed that the fibrocartilage layer thickness was significantly increased, whereas the mature chondrocartilage layer thickness was significantly reduced in β-catenin–activated condyles. Consequently, the ratio of fibrocartilage thickness to total cartilage thickness was significantly elevated compared with controls (Fig. 3F). However, Ki67 staining did not reveal increased proliferative activity in Axin2-lineage cells (Fig. 3E, F), consistent with quantitative analysis showing no significant increase in Ki67-positive cells despite expansion of the fibrocartilage compartment. These findings suggest that sustained β-catenin activation does not drive excessive proliferation but instead perturbs the balance between proliferative maintenance and differentiation. Together, these findings indicate that loss of β-catenin impairs proliferative expansion of fibrocartilage progenitors, leading to defective postnatal condylar growth. These results collectively suggest that canonical Wnt/β-catenin signaling functions as a regulatory axis that maintains the proliferative fibrocartilage progenitor pool while preventing premature chondrogenic differentiation during postnatal mandibular condylar cartilage growth. We therefore next sought to identify transcriptional mediators operating downstream of β-catenin that sustain this proliferative program.

### Foxm1 as a candidate downstream mediator of Wnt/β-catenin signaling

To identify transcriptional mediators associated with canonical Wnt signaling, we performed differential gene expression analysis comparing *H2B-EGFP*–positive and *H2B-EGFP*–negative cells within the MC-progenitor cluster. *Foxm1* emerged as one of the genes significantly enriched in the *H2B-EGFP*–positive population, whereas differentiation-associated genes such as *Tgfb1* were enriched in the *H2B-EGFP*–negative population (Fig. 4A, B). RNAscope in situ hybridization further confirmed that *Foxm1* transcripts were enriched within the fibrocartilage compartment, partially overlapping with Wnt-responsive (*H2B-EGFP*–positive) cells *in vivo*, consistent with preferential expression within the Wnt-active proliferative domain (Extended Data Fig. 6). In contrast, *Igf1* transcripts were predominantly detected in the superficial region of the fibrocartilage layer, indicating spatially restricted expression within the upper fibrocartilage niche (Extended Data Fig. 6). Consistent with these spatial observations, single-cell transcriptomic analysis revealed significantly higher *Foxm1* expression in *H2B-EGFP*–positive cells compared with *H2B-EGFP*–negative cells within the MC-progenitor cluster (Extended Data Fig. 6).

**Figure 4.**
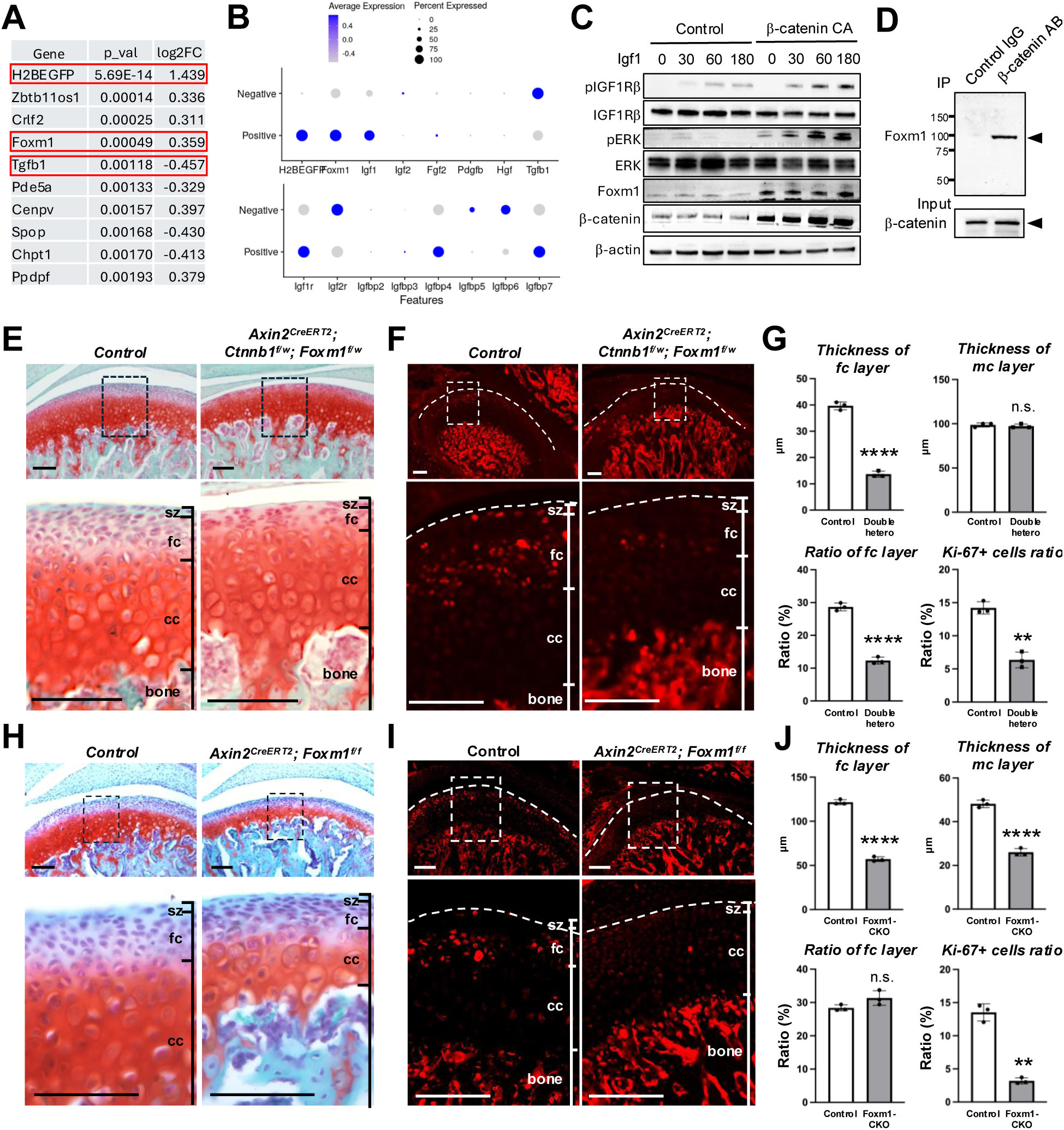
Foxm1 mediates Wnt-associated proliferative programs in condylar fibrocartilage. (A) Differential gene expression analysis was performed comparing *H2B–EGFP*–positive and *H2B–EGFP*–negative cells within the MC-progenitor cluster identified by single-cell RNA sequencing. Genes are ranked by statistical significance. *Foxm1* is significantly enriched in the *H2B–EGFP*–positive population, whereas *Tgfb1* is enriched in the *H2B–EGFP*–negative population, indicating divergent transcriptional programs associated with Wnt activity. (B) Feature plots were generated to visualize expression of representative genes across mandibular condylar cartilage populations. Wnt-responsive cells show enriched expression of *Foxm1* and *IGF* signaling–related genes (*Igf1*, *Igf1r*, *Igfbp4*, *Igfbp7*), whereas Wnt-low populations express *Tgfb1* and related factors (*Igf2r*, *Igfbp5*, *Igfbp6*), supporting distinct signaling states. (C) Western blot analysis was performed in isolated Wnt-responsive cells transfected with control vector or constitutively active β-catenin (S33Y). Cells were stimulated with recombinant IGF1 for the indicated time points. β-catenin activation enhances Foxm1 expression and downstream mitogenic signaling, including ERK and IGF1R phosphorylation, indicating that β-catenin promotes proliferative signaling responses. (D) Co-immunoprecipitation was performed to assess interaction between β-catenin and Foxm1. Cell lysates immunoprecipitated with anti–β-catenin antibody show enrichment of Foxm1 compared with control IgG, indicating a physical association between β-catenin and Foxm1. (E,F) Histological and immunofluorescence analyses were performed on mandibular condyles from control and *Axin2^CreERT2^;Ctnnb1^fl/+^;Foxm1^fl/+^* compound heterozygous mice at P42. H&E staining reveals reduced fibrocartilage thickness, and Ki67 staining shows decreased proliferative activity, indicating cooperative effects of β-catenin and Foxm1 in maintaining fibrocartilage proliferation. Scale bar, 100 μm. (G) Quantification of fibrocartilage thickness and Ki67-positive cells was performed. Compound heterozygous mice show reduced fibrocartilage thickness and decreased proliferation compared with controls. Data are presented as mean ± s.d. Each dot represents one biologically independent animal. (H,I) Histological and immunofluorescence analyses were performed on mandibular condyles from control and *Axin2^CreERT2^;Foxm1^fl/fl^* mice at P42. *Foxm1* deletion results in marked condylar hypoplasia and reduced proliferative activity, indicating a critical role for Foxm1 in fibrocartilage growth. Scale bar, 100 μm. (J) Quantification of cartilage thickness and proliferative indices was performed in Foxm1 conditional knockout mice. *Foxm1* deficiency significantly reduces cartilage growth and proliferation. Data are presented as mean ± s.d. Each dot represents one biologically independent animal. Statistical significance was assessed using two-tailed Student’s t-test. n.s., not significant; **P < 0.01; ****P < 0.0001. Abbreviations: sz, superficial zone; fc, fibrocartilage zone; cc, chondrocartilage zone.

**Figure 5.**
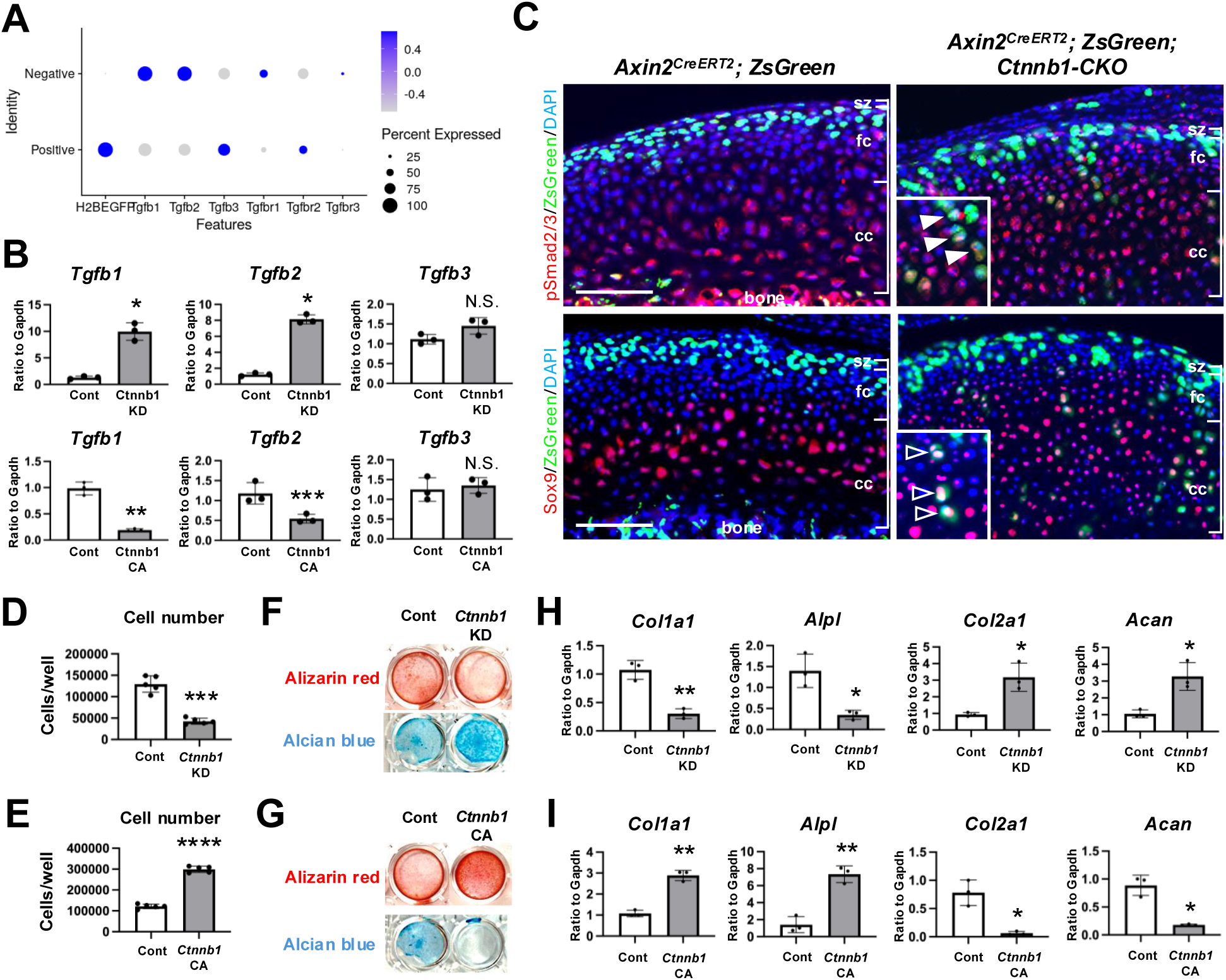
β-catenin restrains TGF-β–Smad signaling to limit chondrogenic commitment. (A) Feature plots were generated to examine expression of TGF-β ligands (*Tgfb1*, *Tgfb2*, *Tgfb3*) and receptors (*Tgfbr1*, *Tgfbr2*) across mandibular condylar cartilage populations. *Tgfb1*, *Tgfb2*, and *Tgfbr1* show inverse correlation with H2B–EGFP expression within the MC-progenitor population, whereas *Tgfb3* and *Tgfbr2* show neutral or positive association, indicating selective antagonism between canonical Wnt activity and specific TGF-β signaling components. (B) Quantitative PCR analysis was performed in cultured Wnt-responsive stem/progenitor cells following Ctnnb1 knockdown (KD) or constitutive β-catenin activation (CA). Ctnnb1 KD increases *Tgfb1* and *Tgfb2* expression, whereas β-catenin CA suppresses their expression, while *Tgfb3* remains unchanged, indicating that β-catenin negatively regulates specific TGF-β ligands. (C) Immunofluorescence staining for phosphorylated Smad2/3 (pSmad2/3) and Sox9 was performed in mandibular condyles from control and *Axin2^CreERT2^;Ctnnb1^fl/fl^* mice. In control mice, pSmad2/3 and Sox9 are largely restricted to the chondrocartilage zone. In contrast, β-catenin–deficient lineage cells show ectopic activation of pSmad2/3 and Sox9 within the fibrocartilage compartment and extension into the chondrocartilage region, indicating enhanced TGF-β–Smad signaling and ectopic activation of chondrogenic programs *in vivo*. Closed arrowheads indicate cells co-expressing pSmad2/3 and ZsGreen, whereas open arrowheads indicate cells co-expressing Sox9 and ZsGreen. Representative images from n = 3 biologically independent animals per genotype are shown. (D,E) Proliferation assays were performed in Wnt-responsive cells following *Ctnnb1* KD or CA. *Ctnnb1* KD reduces cell proliferation, whereas β-catenin CA enhances proliferative activity, indicating that β-catenin positively regulates proliferative capacity. (F) Osteogenic differentiation assays were performed and assessed by Alizarin Red staining. *Ctnnb1* KD reduces mineralized nodule formation, whereas β-catenin CA enhances osteogenic differentiation, indicating promotion of osteogenic programs by β-catenin. (G) Chondrogenic differentiation assays were performed and assessed by Alcian Blue staining. *Ctnnb1* KD increases cartilage matrix deposition, whereas β-catenin CA suppresses chondrogenic differentiation, indicating that β-catenin restrains chondrogenic commitment. (H,I) Quantitative PCR analyses were performed to assess lineage marker expression. *Ctnnb1* KD reduces osteogenic markers (*Col1a1*, *Alpl*) and increases chondrogenic markers (*Col2a1*, *Acan*), whereas β-catenin CA shows the opposite trend, confirming reciprocal regulation of lineage programs. Independent *shRNA* validation experiments are shown in Extended Data Fig. 8. Data are presented as mean ± s.d. from n = 3 biologically independent experiments unless otherwise indicated, each performed in technical triplicate. Statistical significance was assessed using two-tailed Student’s t-test. n.s., not significant; *P < 0.05; **P < 0.01; ***P < 0.001; ****P < 0.0001. Abbreviations: sz, superficial zone; fc, fibrocartilage zone; cc, chondrocartilage zone. Scale bars, 100 μm.

**Figure 6.**
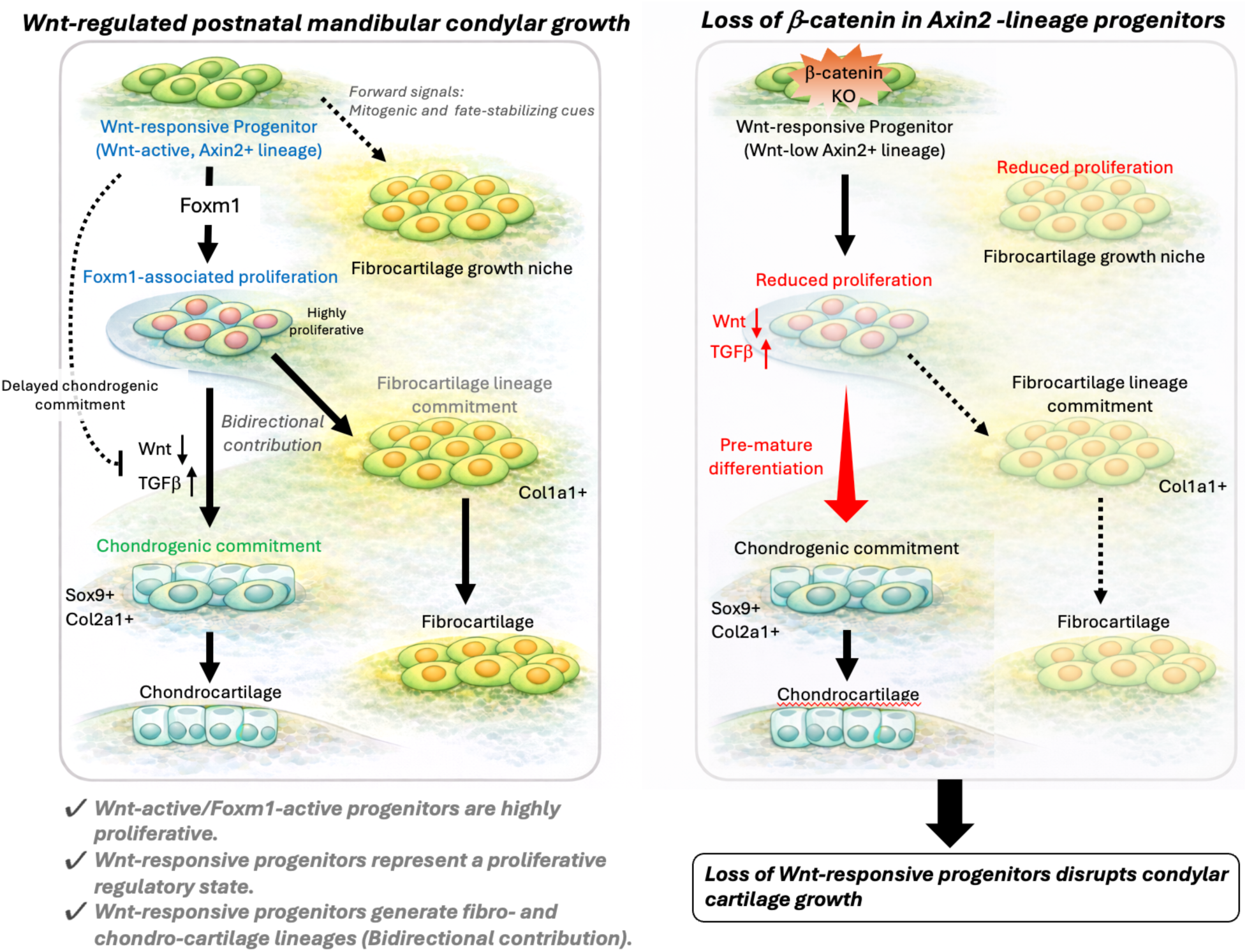
Model for Wnt–Foxm1–dependent regulation of postnatal mandibular condylar cartilage growth. This schematic summarizes a model for how Wnt-responsive fibrocartilage progenitors regulate postnatal mandibular condylar cartilage growth. Axin2-lineage cells reside within the upper fibrocartilage compartment beneath the superficial zone, where a subset of cells exhibits active canonical Wnt signaling and enriched Foxm1 expression. This Wnt/Foxm1-associated state corresponds to the MC-progenitor population identified by single-cell transcriptomic analyses and is characterized by elevated proliferative capacity. Canonical Wnt signaling promotes proliferative expansion of fibrocartilage-associated progenitor cells, at least in part through Foxm1-associated cell-cycle programs, while concurrently restraining TGF-β–Smad–dependent chondrogenic differentiation within this compartment. Lineage tracing demonstrates that Axin2-lineage cells contribute to both fibrocartilage maintenance and chondrocartilage production during postnatal mandibular condylar cartilage growth. Loss of β-catenin disrupts this regulatory balance by reducing the proliferative fibrocartilage compartment and enhancing TGF-β–associated chondrogenic differentiation, ultimately impairing condylar cartilage growth. Together, these findings define a dual regulatory axis in which Wnt/β-catenin signaling coordinates Foxm1-driven proliferative expansion and suppression of TGF-β–dependent chondrogenic differentiation to maintain fibrocartilage progenitor function.

### Foxm1 mediates Wnt-dependent proliferative signaling *in vitro*

To investigate whether Foxm1 functionally contributes to the proliferative program of Wnt-responsive cells, we performed pharmacological and biochemical analyses using isolated Wnt-responsive cells *in vitro*. Pharmacological inhibition of Foxm1 using RCM-1 significantly reduced proliferation of isolated Wnt-responsive cells. Inhibition of canonical Wnt signaling similarly suppressed proliferative activity, and combined inhibition resulted in a further reduction in cell expansion, suggesting that Foxm1 functionally cooperates with β-catenin signaling to promote proliferative capacity (Extended Data Fig. 7). Expression of constitutively active β-catenin (S33Y) increased Foxm1 protein levels (Fig. 4C), supporting a regulatory association between canonical Wnt signaling and Foxm1 expression. However, direct transcriptional regulation of *Foxm1* by β-catenin was not assessed in the present study. To assess mitogenic responsiveness, cells were stimulated with recombinant IGF1 as a defined growth factor input. Under these conditions, β-catenin–activated cells exhibited enhanced ERK phosphorylation compared with control cells (Fig. 4C), indicating increased mitogenic signaling *in vitro*. However, the upstream growth factor pathways responsible for activating this signaling axis *in vivo* remain to be defined. Co-immunoprecipitation experiments demonstrated a physical association between β-catenin and Foxm1 (Fig. 4D). These findings prompted us to examine whether Foxm1 functionally cooperates with β-catenin signaling during postnatal mandibular condylar cartilage growth *in vivo*.

### Genetic interaction between β-catenin and Foxm1 *in vivo*

To evaluate functional interaction between β-catenin and Foxm1 *in vivo*, we generated *Axin2^CreERT2^;Ctnnb1^fl/+^;Foxm1^fl/+^* compound heterozygous mice. At P42, compound heterozygous condyles displayed reduced fibrocartilage thickness and decreased proliferative indices compared with control mice, whereas overall cartilage organization remained largely preserved (Fig. 4E–G). To further assess the requirement for Foxm1 in Axin2-lineage cells, we generated Axin2^CreERT2^;Foxm1^fl/fl^ conditional knockout mice. Conditional deletion of Foxm1 resulted in marked hypoplasia of the mandibular condyle, characterized by thinning of the fibrocartilage layer, reduced numbers of Ki67-positive cells, and decreased overall cartilage thickness (Fig. 4I–K). Collectively, these results demonstrate that Foxm1 functions as a critical mediator associated with canonical Wnt/β-catenin–dependent proliferative expansion and is required for maintenance of postnatal mandibular condylar cartilage growth. These findings prompted us to investigate whether additional signaling pathways downstream of β-catenin regulate the balance between proliferative maintenance and chondrogenic differentiation within Wnt-responsive fibrocartilage cells.

### β-catenin restrains TGF-β–Smad–dependent chondrogenic differentiation

In addition to promoting proliferative expansion through Foxm1-associated mechanisms, canonical Wnt signaling may also regulate lineage progression by modulating TGF-β–dependent chondrogenic differentiation. Notably, *Tgfb1* together with additional pathway components (*Tgfb2*, *Tgfbr1*, *Smad2* and *Smad3*) were enriched in cells with low Wnt reporter activity, suggesting a potential antagonistic relationship between canonical Wnt signaling and TGF-β pathway activation. Immunostaining for phosphorylated Smad2/3 (pSmad2/3) revealed minimal signal within Wnt-responsive cells in the superficial and fibrocartilage zones, whereas strong pSmad2/3 staining was detected in deeper chondrocartilage regions populated by differentiated chondrocytes (Fig. 5C). In *Axin2^CreERT2^;Ctnnb1^fl/fl^;R26^ZsGreen^*mice, β-catenin–deficient lineage cells displayed ectopic extension into the chondrocartilage compartment and exhibited increased pSmad2/3 expression, indicating enhanced TGF-β signaling following β-catenin loss. Consistently, *in vitro* knockdown of *Ctnnb1* upregulated *Tgfb1*, *Tgfb2*, and *Tgfbr1* transcripts, whereas constitutive β-catenin activation suppressed their expression (Fig. 5B). Functionally, *Ctnnb1*-deficient cells exhibited enhanced chondrogenic differentiation, as evidenced by increased Alcian Blue–positive matrix deposition and elevated expression of chondrogenic markers (*Col2a1*, *Acan*) (Fig. 5G, I). Similar results were obtained using an independent shRNA targeting *Ctnnb1*, confirming the reproducibility of the knockdown effects (Extended Data Fig. 8). Conversely, stabilized β-catenin suppressed chondrogenic differentiation under the same conditions. Together, these findings support a model in which canonical Wnt/β-catenin signaling restrains TGF-β–Smad–associated chondrogenic commitment and thereby prevents premature activation of the cartilage differentiation program during postnatal mandibular condyle growth.

Taken together, our analyses reveal that canonical Wnt/β-catenin signaling coordinates two complementary processes within the postnatal mandibular condyle: maintenance of proliferative expansion through Foxm1-associated mechanisms and suppression of TGF-β–Smad–driven chondrogenic differentiation. Loss of β-catenin disrupts both arms of this regulatory axis, resulting in depletion of the fibrocartilage compartment and inappropriate activation of the differentiation program. These findings position Wnt/β-catenin signaling as a central regulatory node that balances proliferative maintenance and lineage progression within Wnt-responsive fibrocartilage progenitors during postnatal mandibular condylar cartilage growth.

## Discussion

In this study, we identify canonical Wnt/β-catenin signaling as a key organizing axis of postnatal MCC growth. Our findings support a model in which Wnt activity coordinates proliferative maintenance within the fibrocartilage compartment while concurrently restraining premature TGF-β–dependent chondrogenic differentiation. By integrating lineage tracing, single-cell transcriptomics, and genetic perturbation, we provide a framework for understanding how proliferative expansion and lineage progression are spatially and temporally balanced in the postnatal mandibular condyle. Together, these findings identify a previously unrecognized Wnt-responsive fibrocartilage progenitor population that orchestrates postnatal mandibular condylar cartilage growth through coordinated control of proliferative maintenance and lineage differentiation.

Canonical Wnt/β-catenin signaling is widely recognized for its roles in progenitor maintenance and lineage specification in multiple tissues (Day et al., 2005; He et al., 1998; Hill et al., 2005; Lowry et al., 2005; Nusse & Clevers, 2017; van de Wetering et al., 2002). In epithelial stem cell niches and skeletal development, Wnt signaling promotes proliferative programs and influences fate decisions. In long bone growth plates, Wnt-responsive resting zone cells contribute to postnatal skeletal growth (Usami et al., 2019). However, the mandibular condyle differs fundamentally from long bone growth plates in both structure and growth mode. MCC growth occurs within a fibrocartilage-dominated environment that must simultaneously support lateral niche expansion and vertical chondrocyte production. How these distinct growth behaviors are coordinated has remained unclear.

Our data identify a Wnt-responsive proliferative population localized predominantly within the fibrocartilage zone. Using complementary reporter systems (*R26-WntVis* and *Axin2^CreERT2^*) and temporally controlled lineage tracing, we demonstrate substantial overlap between active Wnt signaling and Axin2-lineage cells, while acknowledging expected differences due to temporal integration of Cre-based labeling versus real-time reporter activity. Lineage tracing reveals that Axin2-lineage cells expand laterally within fibrocartilage and also generate vertically organized columns of chondrocytes that populate the chondrocartilage compartment. This bidirectional contribution is independently supported by single-cell trajectory inference and RNA velocity analysis. We note that the single-cell transcriptomic dataset was generated from pooled samples derived from two biological replicates but sequenced as a single 10x Genomics library. Although major transcriptional populations were robustly defined, additional independent datasets will be valuable to further validate lineage hierarchies and rare subpopulations. These findings extend and refine previous reports of progenitor-like populations in the mandibular condyle (Acri et al., 2019; Robinson et al., 2015; Ruscitto et al., 2023; Tuwatnawanit et al., 2025).

Rather than defining a rigid unidirectional hierarchy, these findings suggest that Wnt-responsive fibrocartilage cells constitute a dynamic proliferative population capable of supporting both niche expansion and differentiation output. This dual growth behavior provides a conceptual advance in understanding MCC biology: proliferative maintenance and differentiation are not spatially segregated processes but are coordinated within a single Wnt-responsive compartment. In this respect, MCC growth more closely resembles a flexible, context-dependent progenitor system than a classical growth plate–like structure (Chan et al., 2015; Mizuhashi et al., 2018).

Functionally, conditional ablation of β-catenin in Axin2-lineage cells led to depletion of the proliferative fibrocartilage compartment and enhanced chondrogenic differentiation programs. Conversely, constitutive β-catenin activation perturbed compartmental organization without driving hyperproliferation, indicating that physiological levels of Wnt signaling are required to maintain tissue balance. Together, these gain- and loss-of-function analyses demonstrate that β-catenin activity is necessary to sustain proliferative capacity while limiting inappropriate differentiation during postnatal MCC growth (Day et al., 2005; Hill et al., 2005).

At the molecular level, our analyses implicate Foxm1 as an important mediator of Wnt-associated proliferative programs in MCC. Foxm1 expression was enriched within the fibrocartilage zone and positively correlated with Wnt reporter activity in single-cell analyses. Genetic ablation of Foxm1 in Axin2-lineage cells recapitulated key aspects of β-catenin loss, including reduced proliferation and hypoplastic cartilage growth. While these findings support functional cooperation between β-catenin and Foxm1, further studies will be required to determine whether Foxm1 is directly regulated by β-catenin/TCF complexes in this context (Chen et al., 2016; Zhang et al., 2011). Nevertheless, our data position Foxm1 as a critical node linking Wnt signaling to cell cycle control in postnatal MCC.

In parallel, we observed an inverse relationship between Wnt activity and TGF-β–Smad signaling. Wnt-responsive cells exhibited low pSmad2/3 activity, whereas β-catenin ablation resulted in increased TGF-β signaling and enhanced chondrogenic differentiation. These findings are consistent with a model in which canonical Wnt signaling functionally restrains TGF-β–associated chondrogenic commitment, thereby preserving a proliferative fibrocartilage population competent for regulated lineage progression (Guo & Wang, 2009). Thus, Wnt signaling appears to coordinate both proliferative maintenance and differentiation restraint within the same cellular compartment. Importantly, β-catenin deletion reduced proliferation not only within recombined Axin2-lineage cells (ZsGreen-positive) but also in adjacent ZsGreen-negative cells. While ZsGreen negativity does not definitively exclude low-level recombination, the coordinated reduction in proliferation beyond the recombined population is consistent with a non–cell-autonomous component of Wnt signaling in MCC growth regulation. Although the precise paracrine mediators remain to be defined, these findings suggest that Wnt-responsive fibrocartilage cells contribute to establishing a growth-permissive microenvironment. In this regard, Wnt signaling may act not only as a cell-intrinsic regulator but also as an organizer of local niche dynamics.

Several limitations should be acknowledged. *Axin2^CreERT2^*labeling captures a heterogeneous population of Wnt-responsive cells and integrates signaling history over time, and canonical Wnt reporter activity as well as *Axin2* expression are not exclusively restricted to progenitor populations but may also occur in more differentiated fibrocartilage cells. Consequently, although multiple independent approaches support the existence of a Wnt-responsive progenitor population, future studies using more specific lineage markers or clonal tracing strategies will be required to further refine the identity and hierarchy of these cells. Furthermore, although trajectory inference and RNA velocity analyses support a bifurcating hierarchy, definitive lineage relationships will ultimately require complementary approaches such as prospective clonal tracing or independent transcriptomic datasets. Moreover, while genetic evidence supports cooperation between β-catenin and Foxm1, direct transcriptional regulation remains to be established. Future studies employing chromatin profiling and promoter-reporter approaches will further clarify the molecular hierarchy downstream of β-catenin in MCC.

In summary, our study provides a mechanistic framework in which canonical Wnt/β-catenin signaling coordinates proliferative maintenance and differentiation control within the fibrocartilage compartment of the postnatal mandibular condyle. By integrating proliferative expansion through Foxm1-associated mechanisms and restraining TGF-β–dependent chondrogenic commitment, Wnt signaling supports balanced tissue growth and structural integrity during postnatal development. These findings refine current models of TMJ growth and establish Wnt-responsive fibrocartilage cells as key regulators of MCC homeostasis. Collectively, our findings support a revised organizational principle for postnatal mandibular condylar cartilage growth, suggesting that this tissue is maintained not by a classical growth plate–like hierarchy but by a Wnt-regulated fibrocartilage progenitor system that integrates proliferative maintenance and differentiation control within a single spatial niche.

## Materials and Methods

### Mice

To investigate the role of canonical Wnt/β-catenin signaling in regulating postnatal mandibular condylar cartilage growth *in vivo*, we utilized inducible genetic lineage tracing and conditional gene manipulation in mouse models. All animal experiments followed ARRIVE (Animal Research: Reporting of *In Vivo* Experiments) guidelines, and a strict protocol was approved by the Institutional Animal Care and Use Committee of Osaka University Graduate School of Dentistry (Approval No. 04263, 04315). The following mouse lines were used in this study: *Axin2^CreERT2^* mice (purchased from Jackson laboratory, JAX:018867), *Rosa26^ZsGreen^* reporter mice (purchased from Jackson laboratory, JAX mice 007906), *Ctnnb1^fl/fl^* mice (purchased from Jackson laboratory, JAX:004152), *Rosa26^tdTomato^*reporter mice (purchased from Jackson laboratory, JAX mice 007909), *Foxm1^fl/fl^*mice (kindly provided from Dr. Dragana Kopanja) (Wang, Kiyokawa, Dennewitz, & Costa, 2002), *Ctnnb1^exon3-flox^* (kindly provided from Dr. Makoto M. Takeo) (Harada et al., 1999) and *R26-WntVis* reporter mice (purchased from RIKEN BioResource Research Center; CDB0303K https://large.riken.jp/distribution/mutant-list.html) (Takemoto et al., 2016). All mice were maintained on a C57BL6 genetic background. For lineage tracing experiments, *Axin2^CreERT2^* mice were crossed with *Rosa26^ZsGreen^* reporter mice to generate *Axin2^CreERT2^;R26^ZsGreen^*mice. *Rosa26^ZsGreen^* reporter mice were used as a Cre-dependent recombination reporter line. Upon tamoxifen-induced CreERT2 activation, recombined cells irreversibly expressed ZsGreen, allowing identification of cells that underwent Cre-mediated recombination. Recombination efficiency within the fibrocartilage compartment was 23.8 ± 5.0% (mean ± s.d., n = 5 mice), calculated as the proportion of ZsGreen-positive cells among total DAPI-positive nuclei and showing consistent labeling across animals. No ZsGreen signal was detected in control mice lacking tamoxifen administration, confirming minimal Cre leakiness. For conditional deletion of β-catenin in Wnt-responsive cells, *Axin2^CreERT2^* mice were crossed with *Ctnnb1^fl/fl^* mice to generate *Axin2^CreERT2^;Ctnnb1^fl/fl^*mice. For conditional deletion of *Foxm1* in Wnt-responsive cells, *Axin2^CreERT2^*mice were crossed with *Foxm1^fl/fl^* mice to generate *Axin2^CreERT2^;Foxm1^fl/fl^*mice. For compound heterozygous analyses, *Axin2^CreERT2^* mice were crossed with *Ctnnb1^fl/+^;Foxm1^fl/+^* mice.

For constitutive activation of β-catenin, *Axin2^CreERT2^*mice were crossed with *Ctnnb1^exon3-flox^* mice to generate *Axin2^CreERT2^;Ctnnb1^exon3-flox^*mice, in which Cre-mediated recombination stabilizes β-catenin by deletion of exon 3. For compound heterozygous analyses, *Axin2^CreERT2^* mice were crossed with *Ctnnb1^fl/+^;Foxm1^fl/+^* mice to generate *Axin2^CreERT2^;Ctnnb1^fl/+^;Foxm1^fl/+^* mice. For combined lineage tracing and β-catenin deletion, *Axin2^CreERT2^;Ctnnb1^fl/fl^* mice were crossed with *Ctnnb1^fl/fl^;R26^ZsGreen^*mice. For visualization of canonical Wnt/β-catenin signaling activity, *R26-WntVis* reporter mice were used, in which H2B–EGFP expression is driven by TCF/LEF-responsive elements. For combined lineage tracing and Wnt activity analysis, *Axin2^CreERT2^* mice were crossed with *Rosa26^tdTomato^*and *R26-WntVis* reporter mice to generate *Axin2^CreERT2^;R26^tdTomato^;R26-WntVis* mice. In this model, Tomato fluorescence marks Axin2-lineage cells following tamoxifen-induced recombination, whereas H2B–EGFP expression indicates active canonical Wnt/β-catenin signaling. All compound mutant and control littermates were generated from the same breeding scheme and analyzed in parallel. Unless otherwise stated, female mice were used due to reduced fertility observed in male *Axin2^CreERT2^* mice. Control mice included both *Cre*-negative littermates and *Axin2^CreERT2^*-positive mice lacking *floxed* alleles, as appropriate for each experiment. All control animals were administered tamoxifen using the same protocols as experimental groups to control for potential effects of Cre activation and tamoxifen treatment. Mice were housed under specific pathogen–free conditions with controlled temperature and humidity, a 12-h light/12-h dark cycle (lights on from 8 to 20), and ad libitum access to food and water.

### Single-cell RNA sequencing, trajectory inference, and RNA velocity analysis

Mandibular condylar tissues were collected at P16, a stage characterized by active postnatal condylar growth and expansion of the fibrocartilage compartment, thereby allowing capture of proliferative Wnt-responsive progenitor populations during ongoing tissue growth. Cells from two biological replicates were pooled prior to library preparation and sequenced as a single 10x Genomics library. Single-cell RNA-sequencing libraries were generated using the Chromium Next GEM Single Cell 3′ Reagent Kits v3.1 (10x Genomics) according to the manufacturer’s instructions. Library preparation and sequencing were performed at the Center for Medical Innovation and Translational Research (CoMIT), Osaka University. Libraries were sequenced on an Illumina NovaSeq 6000 platform using paired-end sequencing (28 bp for read 1 and 91 bp for read 2). In total, 766,634,800 reads were generated, corresponding to 45.6 Gbp of sequence data. The overall GC content was 46.95%, with Q20 and Q30 scores of 94.77% and 89.02%, respectively. Raw sequencing data were demultiplexed using bcl2fastq (v2.20.0). Sequencing quality was evaluated using FastQC (v0.11.7). Alignment to the mouse reference genome, barcode processing, and gene counting were performed using Cell Ranger (v7.0.1) with default parameters. Downstream analyses were conducted in R (v4.2.3) using the Seurat package. The percentage of mitochondrial transcripts was calculated using the PercentageFeatureSet function (pattern = “^mt-”). Cells with fewer than 200 detected genes, more than 8,000 detected genes, or greater than 20% mitochondrial gene content were excluded. Putative doublets were excluded based on abnormally high gene counts relative to the overall distribution of detected genes across cells. Because all cells were derived from a single 10x Genomics library, batch integration was not required. Gene expression values were normalized using the LogNormalize method with a scale factor of 10,000. Highly variable genes were identified using the “vst” method (2,000 features). The data were subsequently scaled and subjected to principal component analysis (PCA). Significant principal components were selected based on JackStraw analysis and ElbowPlot inspection. Cells were clustered using the shared nearest neighbor (SNN) modularity optimization algorithm with a resolution of 0.5 and visualized using Uniform Manifold Approximation and Projection (UMAP). To evaluate whether cluster identities were driven by cell-cycle heterogeneity, cell-cycle regression was performed during data scaling using canonical S and G2/M gene sets. The MC-progenitor cluster remained identifiable after cell-cycle regression, indicating that this population was not solely defined by cell-cycle–associated transcriptional programs. Based on marker gene expression, mandibular condylar cells were broadly classified into fibrocartilage, mature chondrocyte, and synovial populations. Cells expressing high levels of Col1a1 were designated as flattened chondrocytes and further subdivided into three subclusters (clusters 0–2) based on differential expression of Cdh11, Tspan2, and Irf1. Clusters expressing high levels of Col2a1 were defined as mature chondrocytes and subdivided into two subclusters (clusters 3 and 4) based on Folr2 and M1ap expression. A cluster enriched for Col10a1 was classified as hypertrophic chondrocytes (cluster 6). Cells expressing Prg4 were defined as superficial zone cells (cluster 7), whereas cells enriched for Pthlh were classified as polymorphic cells (cluster 8). To infer developmental trajectories and lineage relationships among mandibular condylar cell populations, pseudotime trajectory analysis was performed using Monocle3 (v1.3.1). Cells were ordered in pseudotime based on transcriptional similarity, and the trajectory root was defined at the MC-progenitor cluster based on enrichment of Wnt-responsive (EGFP-positive) cells and its position at the origin of the inferred trajectory. To independently evaluate lineage directionality, Velocity analysis was performed using the stochastic/dynamical model implemented in scVelo following the standard workflow (Bergen, Lange, Peidli, Wolf, & Theis, 2020). Spliced and unspliced transcript counts were extracted from aligned BAM files using Velocyto (v0.17.17). Velocity estimation was performed using the dynamical model implemented in scVelo following the standard preprocessing workflow. Genes were filtered based on a minimum shared count threshold (min_shared_counts = 20) and normalized using scVelo’s default normalization procedure. First- and second-order moments were computed using the top 30 principal components and 30 nearest neighbors. Velocity vectors were projected onto the UMAP embedding generated from the Seurat analysis to visualize predicted transcriptional state transitions. Wnt-responsive cells were defined based on detectable expression of the H2B-EGFP transcript above background levels in the normalized expression matrix. Differentially expressed genes (DEGs) were identified in clusters enriched for EGFP-positive (Wnt-responsive) cells. Functional enrichment analysis of DEGs was conducted using the KEGG pathway database to identify cell-cycle-related genes and signaling components potentially associated with β-catenin–dependent transcriptional programs.

### Tamoxifen administration and experimental design

Tamoxifen (Sigma-Aldrich) was dissolved in corn oil (Sigma-Aldrich) at a concentration of 10 mg/mL and administered by intraperitoneal injection at a dose of 40 mg/kg body weight at postnatal day 14 (P14). For lineage tracing analyses, *Axin2^CreERT2^;ZsGreen* mice were harvested 2, 7, or 28 days after tamoxifen administration. For conditional knockout and gain-of-function analyses, *Axin2^CreERT2^;Ctnnb1^fl/fl^*, *Axin2^CreERT2^;Ctnnb1^fl/wt^;Foxm1^fl/wt^*, *Axin2^CreERT2^;Foxm1^fl/fl^*, and *Axin2^CreERT2^;Ctnnb1^exon3-flox^* female mice were harvested 7 or 28 days after tamoxifen administration. For experiments involving multiple tamoxifen administrations, tamoxifen was administered intraperitoneally once daily for five consecutive days starting at P14, and mice were harvested 28 days after the initial tamoxifen injection. Control mice, which expressed *Axin2^CreERT2^* alone without floxed alleles, were also administered tamoxifen following the same injection protocols to control for potential effects of tamoxifen and CreERT2 activation. Experimental group sizes (n) represent biologically independent animals unless otherwise stated.

### Tissue preparation and cryosectioning

Mice were euthanized by carbon dioxide inhalation, and craniofacial tissues were immediately harvested. Specimens were fixed in 4 % paraformaldehyde in 0.1 M phosphate-buffered saline (PBS; pH 7.4) at 4 °C for 24 h. For decalcified sections, samples were decalcified in Osteosoft® (101728, Millipore) at 4 °C for 2 weeks, with decalcification solution replaced every 3 days. After decalcification, tissues were cryoprotected by sequential immersion in 10 %, 20 %, and 30 % sucrose solutions in PBS at 4 °C, until fully infiltrated. Samples were embedded in Tissue-Tek OCT compound (Sakura Finetek, Japan) and rapidly frozen on dry ice. Sagittal cryosections were cut at a thickness of 15 µm using a cryostat (CM1950; Leica Microsystems, Wetzlar, Germany). Sections were stored at −30 °C until further use. For non-decalcified frozen sections used for in situ hybridization, tissues were processed using the Kawamoto film method as previously described (Inubushi, Lemire, Irie, & Yamaguchi, 2018), and cryosections were cut at a thickness of 14 µm.

### Immunofluorescence staining

Immunostaining of frozen sections was performed as previously described (Inubushi, Nozawa, Matsumoto, Irie, & Yamaguchi, 2017). Cryosections were air-dried and rehydrated in PBS. Sections were blocked with 5% BSA and 0.3% Triton-X100 in PBS for 60 minutes at room temperature. Primary antibodies were diluted in blocking solution and incubated overnight at 4 °C. The following primary antibodies were used: anti–Ki67 (Cat No. ab16667, Abcam), anti–Sox9 (Cat No. 82630, Cell Signaling), anti–phospho Smad2/3 (Cat No. AP0548, ABClonal). After washing in PBS, sections were incubated with species-appropriate fluorophore-conjugated secondary antibodies donkey anti-rabbit IgG Alexa Fluor 555 conjugate (Invitrogen, S32355, 1:500) and DAPI (Dojindo, D523, 1:250) for 1 hour at room temperature. Sections were mounted using aqueous fluorescence mounting medium (DAKO). Images were acquired using all-in-one fluorescence microscope (Keyence BZ-X710) with identical acquisition settings for all experimental groups.

### Image quantification

Quantitative analyses were performed using Image J software (version 6). Fibrocartilage and chondrocartilage thickness were measured on sagittal sections spanning the central region of the mandibular condyle, defined as the midpoint between the medial and lateral poles of the condyle. Fibrocartilage ratio is calculated as percentage of total thickness (Fibrocartilage and chondrocartilage). For each mouse, 3 independent non-adjacent sections were analyzed and averaged. Ki67-positive cells were quantified as percentage of total nuclei in the fibrocartilage layer of the section. All analyses were performed by investigators blinded to genotype.

### Micro–computed tomography (micro-CT) analysis

Mandibular condyles were scanned using a micro-CT system (R_mCT2, Rigaku). Scanning was performed at 90 kV and 200 μA with a voxel size of 10 μm. Three-dimensional reconstruction was performed using VGStudio. Condylar length was measured using reconstructed images based on anatomical landmarks.

### EdU label-retention assay

EdU (5-ethynyl-2′-deoxyuridine; Sigma-Aldrich; catalog no. 900584) was administered to mice by intraperitoneal injection at a dose of 10 mg/kg body weight. For EdU label-retention experiments, mice were harvested 28 days after EdU administration to identify slow-cycling cells. EdU incorporation was detected on frozen sections using the Click-iT™ EdU Imaging Kit (Thermo Fisher Scientific) according to the manufacturer’s instructions. Sections were counterstained with DAPI and imaged by fluorescence microscopy.

### RNAscope in situ hybridization

RNAscope *in situ* hybridization was performed using the RNAscope Multiplex Fluorescent Reagent Kit v2 (Advanced Cell Diagnostics, ACD) according to the manufacturer’s instructions. Briefly, cryosections were pretreated following the standard ACD protocol and hybridized with RNAscope probes targeting Foxm1 mRNA (ACD; catalog no. 503481), IGF1 (ACD; catalog no. 443901-C3). As a negative control, sections were hybridized with a negative control probe (ACD; catalog no. 310043). Signal amplification was performed using the RNAscope Multiplex Fluorescent v2 amplification system. Detection was carried out using TSA Plus Cyanine 3 (Akoya Biosciences) as the fluorophore. Nuclei were counterstained with DAPI (Dojindo, Japan). Sections were mounted using Dako Fluorescence Mounting Medium (Agilent Technologies) and imaged using a BZ-X700 fluorescence microscope (Keyence, Japan).

### Isolation and culture of mandibular condylar cells

Mandibular condylar cartilage was dissected from P14 mice and enzymatically digested with 24 mg/mL collagenase type II (Worthington Biochemical Corporation, USA) in phosphate-buffered saline (PBS) at 37 °C for 30 minutes with gentle agitation. Dissociated cells were passed through a 14-µm cell strainer to obtain a single-cell suspension and cultured in stem cell medium (MesenCult™ Expansion Kit, STEMCELL Technologies) under standard conditions. After expansion, EGFP-positive cells were isolated by fluorescence-activated cell sorting (FACS) using a SH800S Cell Sorter (SONY, Japan). Cell sorting gates were defined based on forward scatter (FSC) and side scatter (SSC) to exclude debris and dead cells, followed by gating for singlets and EGFP-positive cells based on fluorescence intensity compared to non-reporter control samples. Sorted cells were subsequently transduced with a temperature-sensitive SV40 large T antigen–expressing adenoviral vector (VectorBuilder, USA) at a multiplicity of infection (MOI) of 10 and maintained at 33 °C to allow conditional cell expansion. For surface marker analysis, cells were incubated with fluorophore-conjugated antibodies against CD29, Sca-1, CD44, CD105, and CD106 as positive markers, and CD11 and CD45 as negative markers, for 30 minutes at room temperature. After washing, cells were analyzed using SH800S analysis software (SONY, Japan).

### Adenoviral-mediated β-catenin manipulation

Wnt-responsive cells were infected with adenoviral vectors purchased from VectorBuilder (China). For β-catenin knockdown, two independent adenoviruses expressing short hairpin RNAs targeting distinct sequences of mouse Ctnnb1 (pAV[shRNA-EGFP-U6]-mCtnnb1-shRNAs) were used. For β-catenin gain-of-function experiments, cells were infected with pAV[Exp-EGFP-CMV]-Human Ctnnb1(S33Y)-FLAG, which encodes a constitutively active form of human β-catenin. Control cells were infected with a scrambled shRNA-expressing adenovirus (pAV[shRNA-EGFP-U6]-Scramble-shRNA) or an empty control vector (pAV[Exp-EGFP-CMV]). All adenoviral vectors were generated using the same backbone (VectorBuilder), and infections were performed under identical conditions (MOI = 5, same viral lot, and identical culture duration). This experimental design enabled direct comparisons between β-catenin loss- and gain-of-function conditions relative to matched controls. After infection, cells were cultured for 72 h in low-glucose DMEM supplemented with 10% fetal bovine serum and 1% penicillin–streptomycin at 37 °C in a humidified atmosphere containing 5% CO₂.

### Proliferation assays

Wnt-responsive cells were isolated from mandibular condylar cartilage of R26-WntVis mice at P16 and plated at a density of 1 × 10⁴ cells per 3.5-cm culture dish (AGC Techno glass, Japan). Cells were cultured in low glucose DMEM supplemented with 10% fetal bovine serum (FBS) (FUJIFILM Wako Pure Chemical Corporation, Japan). Pharmacological treatments were performed using the Foxm1 inhibitor RCM-1, the β-catenin inhibitor XAV-939, or a canonical Wnt pathway activator (Wnt agonist 1) (all from Selleck Chemicals, Japan). Each compound was dissolved in dimethyl sulfoxide (DMSO) and added to the culture medium at a final dilution of 1:200. Final working concentrations were 4 µM or 20 µM for RCM-1, 1 µM or 5 µM for XAV-939, and 1 nM for Wnt agonist 1. Control cells were treated with an equivalent concentration of DMSO (1:200 dilution). Cells were cultured under treatment conditions for 7 days, after which cell numbers were quantified using a Countess™ 3 Automated Cell Counter (Thermo Fisher Scientific). Each experiment was performed with biologically independent replicates.

### *In vitro* differentiation assays

Sorted cells were seeded at a density of 5 × 10⁴ cells per well in 24-well plates (AGC Techno glass, Japan) and cultured under standard conditions. *In vitro* trigenic differentiation assays were performed as previously described (Inubushi et al., 2017). For osteogenic differentiation, cells were cultured in osteogenic induction medium consisting of 10% fetal bovine serum (FBS), 10 mM β-glycerophosphate, and 50 μg/mL L-ascorbic acid, as previously described. For adipogenic differentiation, cells were cultured in adipogenic induction medium containing 10% FBS, 200 μM indomethacin, 10 μg/mL insulin, 0.5 mM 3-isobutyl-1-methylxanthine (IBMX), and 1 μM dexamethasone. For chondrogenic differentiation, DMEM containing 100 nM dexamethasone, 50 μg/ml ascorbic acid-2-phosphate, 100 μg/ml sodium pyruvate, 40 μg/ml L-proline, 1× ITS+, and 10 ng/ml recombinant mouse TGF-β1 (Cat No. 7666-MB-005). Differentiation was assessed after 3 weeks by Alizarin Red S staining for osteogenic differentiation, Oil Red O staining for adipogenic differentiation, and Alcian Blue staining for chondrogenic differentiation, following standard protocols. Stained cultures were imaged, and representative fields are shown. Quantification was not performed.

### RNA extraction and qPCR analysis

β-catenin–constitutively activated cells (S33Y-transfected), *Ctnnb1* knockdown cells transfected with two independent shRNAs targeting Ctnnb1 (*shCtnnb1* and *shCtnnb1_2*), and matched control (Control and *shControl*) cells were seeded at a density of 1 × 10⁵ cells per well in 24-well culture plates (AGC Techno glass, Japan). After 3 weeks of incubation under osteogenic or chondrogenic differentiation conditions, total RNA was extracted using the RNeasy Mini Kit (Qiagen, Germany) and reverse-transcribed into cDNA using oligo(dT) primers and reverse transcriptase (Takara Bio, Japan), as previously described (Inubushi et al., 2012). Quantitative real-time PCR was performed using SYBR Green PCR Master Mix (Takara Bio, Japan) on a StepOne Real-Time PCR System with StepOne Software v2.1 (Applied Biosystems). Expression levels of Col1a1, Alp, Col2a1, Acan, and Ctnnb1 were normalized to β-actin (Actb), which was confirmed to be stably expressed across experimental conditions. Primer sequences are provided in Extended Data.

### Immunoprecipitation

Co-immunoprecipitation was performed using the Dynabeads™ Co-Immunoprecipitation Kit (Cat No. 14321D, Thermo Fisher Scientific) according to the manufacturer’s instructions. Briefly, Wnt-responsive stem/progenitor cells were lysed in ice-cold lysis buffer containing 1% NP-40, 25 mM HEPES (pH 7.5), 150 mM NaCl, 5 mM MgCl₂, and a protease inhibitor cocktail (Cat No. 3969-21, Nacalai)and phosphatase inhibitor cocktail (Cat No. 7575-51, Nacalai). Lysates were clarified by centrifugation at 14,000 × g for 15 min at 4 °C. For immunoprecipitation, clarified lysates were incubated with either anti–β-catenin antibody (Cat No. 8480, Cell Signaling) or isotype-matched control IgG at 4 °C with gentle rotation for overnight incubation. Antibody–protein complexes were captured using Dynabeads magnetic beads, followed by multiple washes with the provided wash buffer to remove nonspecific binding. Immunoprecipitated proteins were eluted using the supplied elution buffer and subjected to immunoblotting.

### Immunoblotting

Wnt-responsive stem/progenitor cells were transfected with either control vector or constitutively active β-catenin (S33Y). Cells were stimulated with recombinant mouse IGF1 (Cat No. 791-MG-050, R&D systems) for 0, 30, 60, or 180 min. Cells were then immediately lysed and processed for immunoblotting. For immunoblot analysis, total cell lysates or immunoprecipitated samples were resolved by SDS–PAGE on 8–16% Tris–glycine gels (Thermo Fisher Scientific) and transferred onto PVDF membranes (Immobilon-P, Millipore). Membranes were blocked with 5% non-fat dry milk or 5% BSA in Tris-buffered saline containing 0.1% Tween-20 (TBST) and incubated with primary antibodies overnight at 4 °C. The following primary antibodies were used: anti-Foxm1 (Cat No. 13147-1-AP, Proteintech), anti–β-catenin (Cat No. 8480, Cell Signaling), anti–phospho-ERK1/2 (Thr202/Tyr204) (Cat No. 5726, Cell Signaling), anti-ERK1/2 (Cat No. 4695, Cell Signaling), anti–phospho-IGF1R (Cat No. 3024, Cell Signaling), anti-IGF1R (Cat No. 3027, Cell Signaling). After washing, membranes were incubated with Goat anti-mouse IgG (H + L)-horseradish peroxidase–conjugated secondary antibody (Cat No. 1706516, Bio-Rad) or Goat anti-rabbit IgG (H + L)-horseradish peroxidase–conjugated secondary antibody (Cat No. 1706515, Bio-Rad) for 1 h at room temperature. Signals were detected using enhanced chemiluminescence (ECL) reagents (Chemi-Lumi One, Nacalai, Japan) and visualized with a chemiluminescence imaging system (iBright FL 1500 imaging system, Thermo Fisher Scientific).

### Statistical analysis

Biologically independent replicates were defined as samples derived from different animals for *in vivo* experiments and from independent cell isolations for *in vitro* assays, unless otherwise stated. No statistical methods were used to predetermine sample size. Statistical analyses were performed using GraphPad Prism 8 (GraphPad Software, USA). Data distribution was assessed prior to analysis to confirm the appropriateness of parametric tests. Comparisons between two groups were performed using two-tailed Student’s t-test. Comparisons among multiple groups were performed using one-way ANOVA followed by Tukey’s post hoc test, or two-way ANOVA followed by Šidák’s post hoc test, as appropriate. A p value < 0.05 was considered statistically significant. Data are presented as mean ± s.e.m., unless otherwise stated. Sample sizes (n) and statistical details are provided in the figure legends. Quantitative analyses were performed using biologically independent animals, cells, or samples, as indicated in the figure legends. Immunostaining experiments were repeated independently at least three times with similar results. No randomization was used to allocate experimental groups, and no animals were excluded from the analyses. Investigators were blinded to genotype during image quantification.

## Ethics Statement

All animal experiments were approved by the Institutional Animal Care and Use Committee of Osaka University Graduate School of Dentistry (approval numbers 04263 and 04315) and were conducted in accordance with the ARRIVE guidelines and institutional regulations for animal welfare. No data were excluded from the analyses. No wild animals or field-collected samples were used in this study.

## Data availability

All data supporting the findings of this study are available within the paper and its Supplementary Information. The single-cell RNA-seq dataset generated in this study will be made available upon publication. All numerical source data underlying the graphs and charts in the main and supplementary figures are provided as Supplementary Data files. Additional raw data and analysis files are available from the corresponding author upon reasonable request.

## Acknowledgments

We thank Ms. Yuki Okamoto, Ms. Satomi Gion, and Ms. Mayumi Yoshimoto for excellent animal care and for technical assistance with histological, molecular, and protein analyses. We are grateful to Dr. Dragana Kopanja for providing the *Foxm1^f/f^* mice and to Prof. Makoto Taketo for providing the *Ctnnb1^exon3-flox^*mice.

## Funding

This work was supported by Grants-in-Aid for Scientific Research (24K02652 to T.I., 24K23619 to R.K.) from the Japan Society for the Promotion of Science and by the Japan Science and Technology Agency FOREST Program (JPMJFR220J to T.I.).

## Declaration of Interests

The authors declare no competing interests.

## Author Contributions

R. Kani and Y. Tanida contributed to data acquisition, analysis, and interpretation, drafted the manuscript; T. Inubushi, contributed to conception and design, data acquisition, analysis, and interpretation, drafted and critically revised the manuscript; Y. Usami, T. Iwayama, W. Deyang, J. Ye, S. Kusano and JI. Sasaki contributed to data acquisition, and analysis; Y. Shiraishi, H. Kurosaka, D. Kopanja, M. Takedachi, S. Murakami and T. Yamashiro critically revised the manuscript; All authors gave final approval and agreed to be accountable for all aspects of the work.

---

**Extended Data Fig. 1.**
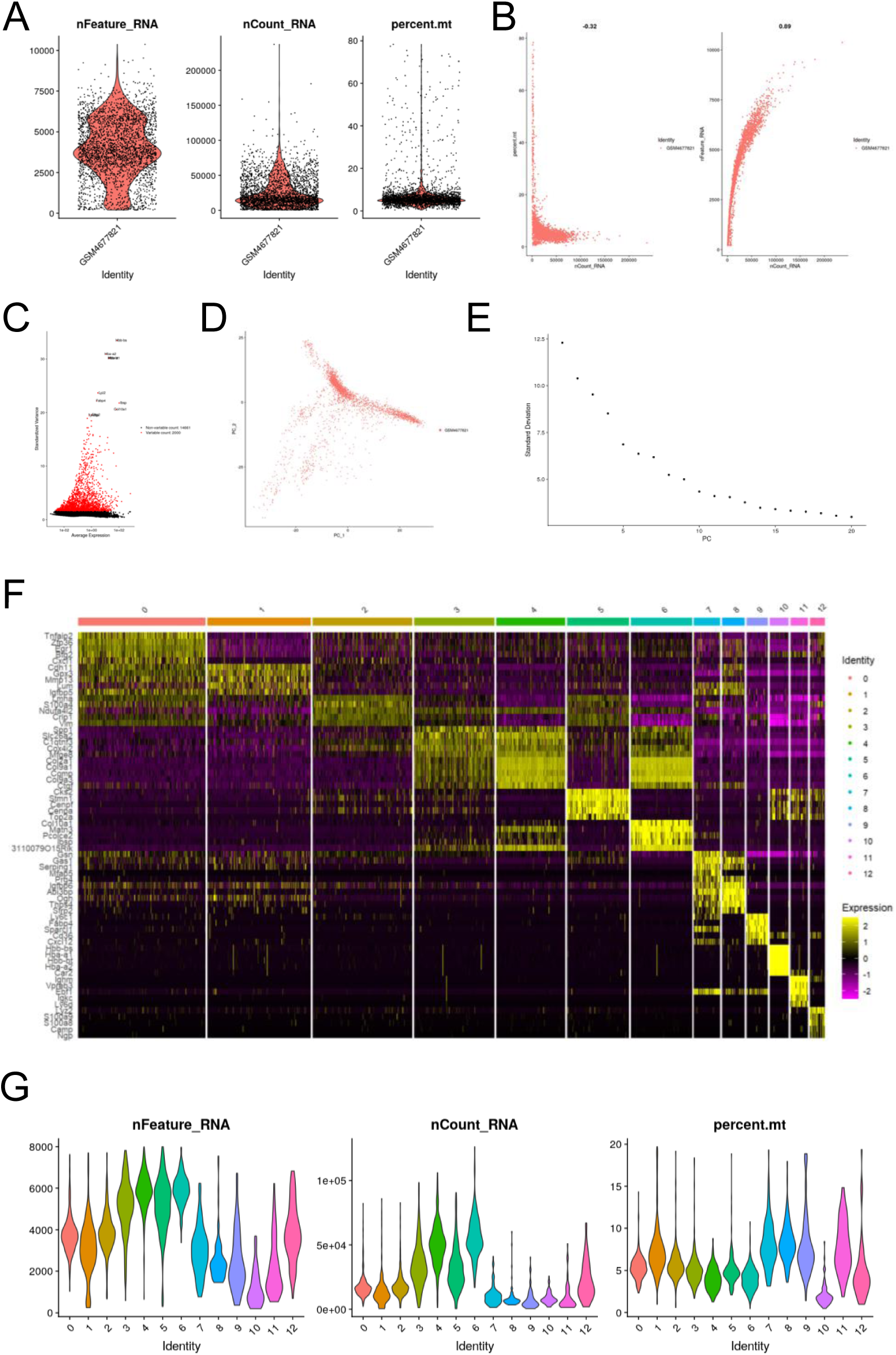
Quality control and preprocessing of single-cell RNA sequencing data. (A) Violin plots showing the distribution of detected genes (nFeature_RNA), total transcript counts (nCount_RNA), and mitochondrial transcript percentage (percent.mt) across cells prior to filtering. (B) Scatter plots showing relationships between nCount_RNA and percent.mt (left) and between nCount_RNA and nFeature_RNA (right). (C) Identification of highly variable genes used for downstream clustering. (D) Principal component analysis (PCA) of single-cell transcriptomes. (E) Elbow plot used to determine the number of principal components included in downstream analyses. (F) Heatmap of the top differentially expressed genes across clusters. (G) Violin plots showing distributions of nFeature_RNA, nCount_RNA, and percent.mt across clusters, confirming comparable RNA complexity and mitochondrial transcript content among clusters.

**Extended Data Fig. 2.**
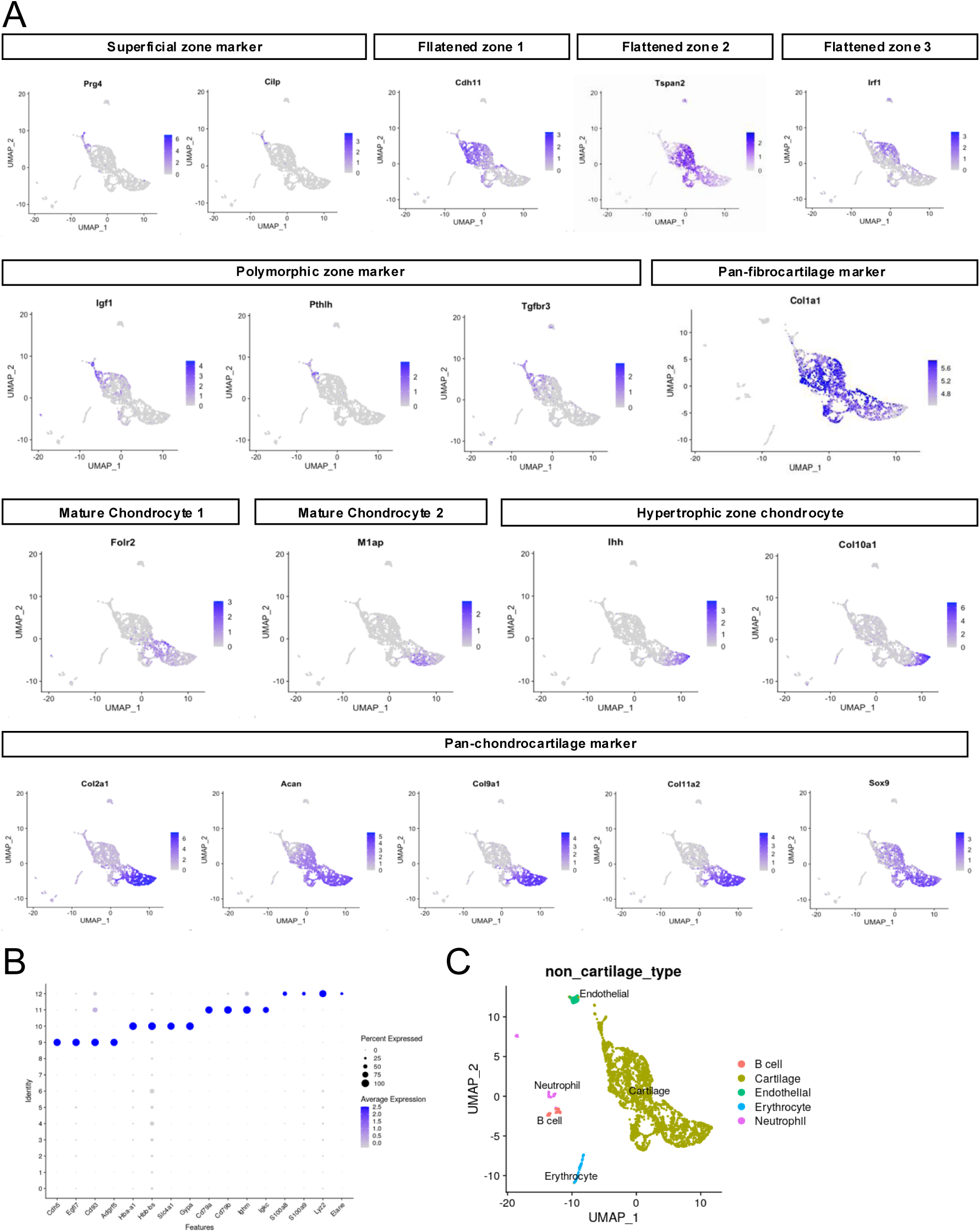
Marker gene expression defining fibrocartilage and chondrocartilage cell populations. (A) Feature plots showing expression of representative marker genes used to annotate fibrocartilage (*Prg4, Cilp, Cdh11, Tspan2, Irf1*), chondrocartilage (*Col2a1, Acan, Col9a1, Col11a2, Sox9*), and hypertrophic chondrocytes (*Col10a1*). (B) Dot plot summarizing expression of canonical lineage markers across clusters. (C) UMAP visualization highlighting non-cartilage cell populations including endothelial cells (*Cdh5*), erythrocytes (*Hbb/Hba*), B cells (*Cd79a/Cd79b*), and neutrophils (*S100a8/S100a9*).

**Extended Data Fig. 3.**
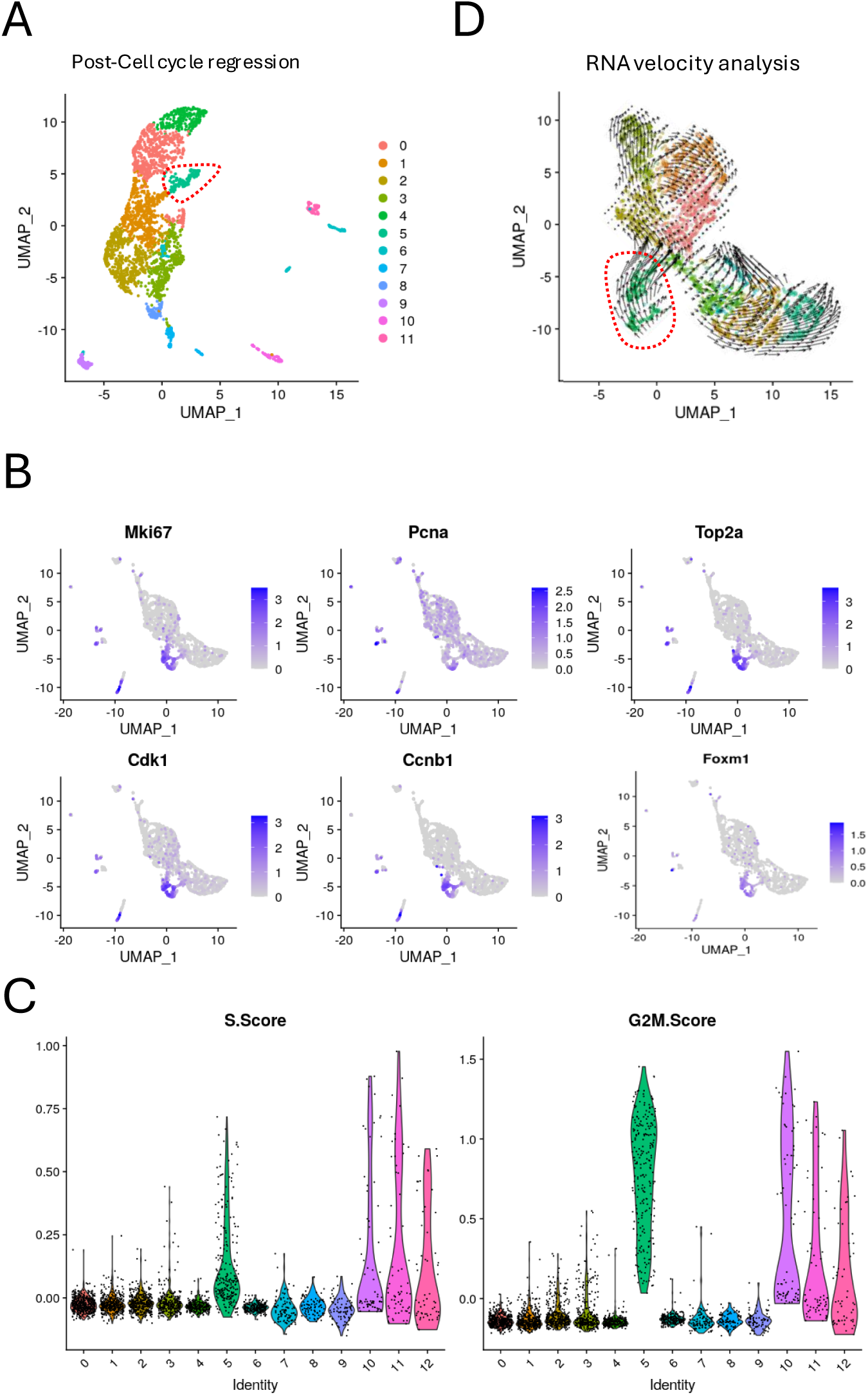
Cell-cycle regression and RNA velocity analysis of condylar cartilage populations. (A) UMAP visualization after regression of cell-cycle effects showing that the MC-progenitor cluster remains preserved following cell-cycle correction. (B) Feature plots showing expression of representative cell-cycle–associated genes (*Mki67, Pcna, Top2a, Cdk1, Ccnb1*) and Foxm1. (C) Violin plots showing S phase and G2/M phase cell-cycle scores across clusters. (D) RNA velocity analysis using scVelo showing directional transcriptional flows from the MC-progenitor cluster toward fibrocartilage and chondrocartilage populations.

**Extended Data Fig. 4.**
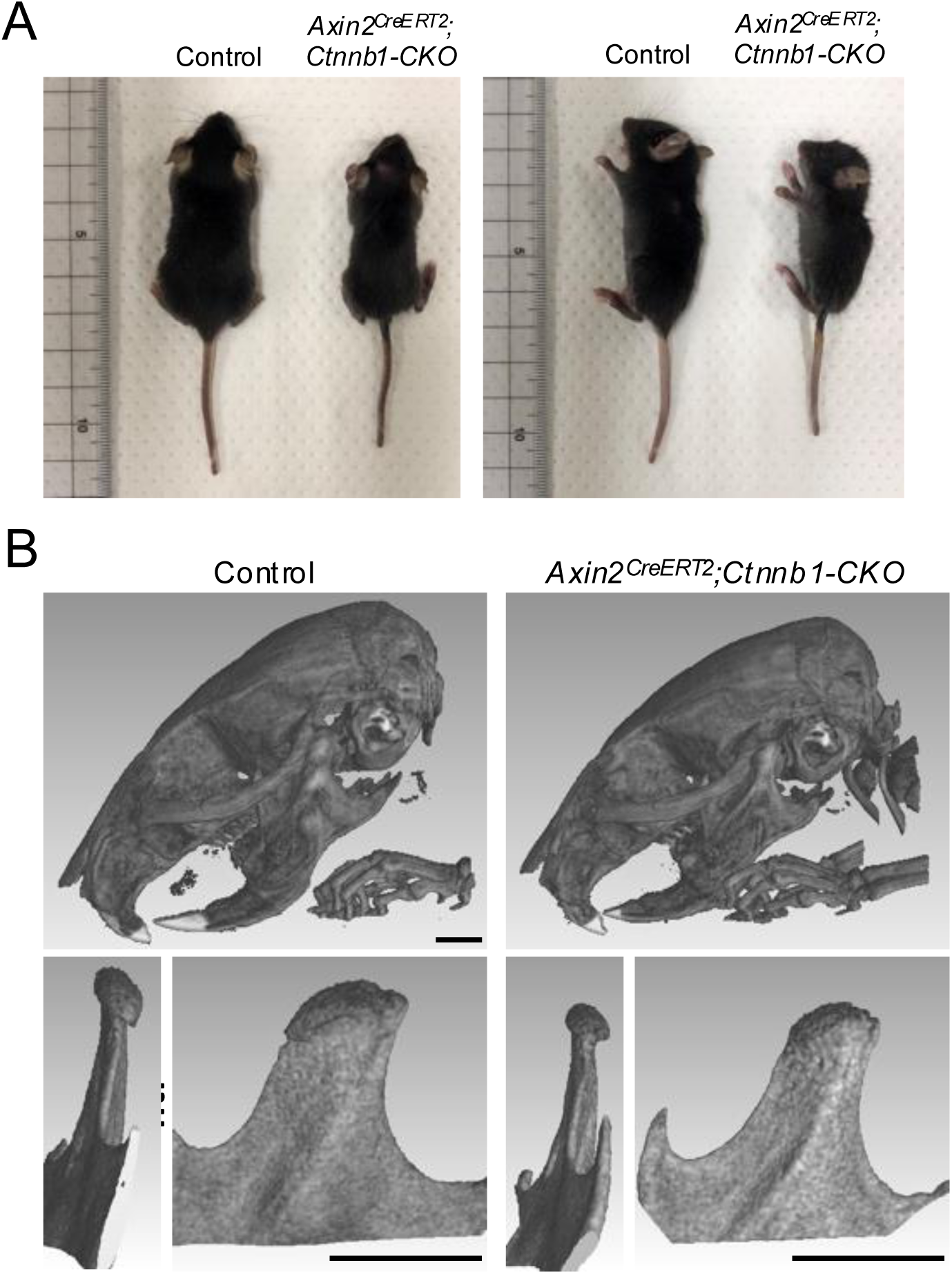
Gross morphology and micro-CT analysis of mandibular condyle in β-catenin–deficient mice. (A) Gross morphology of control and *Axin2^CreERT2^;Ctnnb1^fl/fl^* mice at P42. Representative images from dorsal (top) and lateral (side) views show no overt differences in overall body size or morphology between genotypes. (B) Micro-CT analysis of mandibular condyles from control and *Axin2^CreERT2^;Ctnnb1^fl/fl^* mice at P42. Three-dimensional reconstructions reveal reduced condylar length in β-catenin–deficient mice compared with littermate controls. Scale bars: 2 mm.

**Extended Data Fig. 5.**
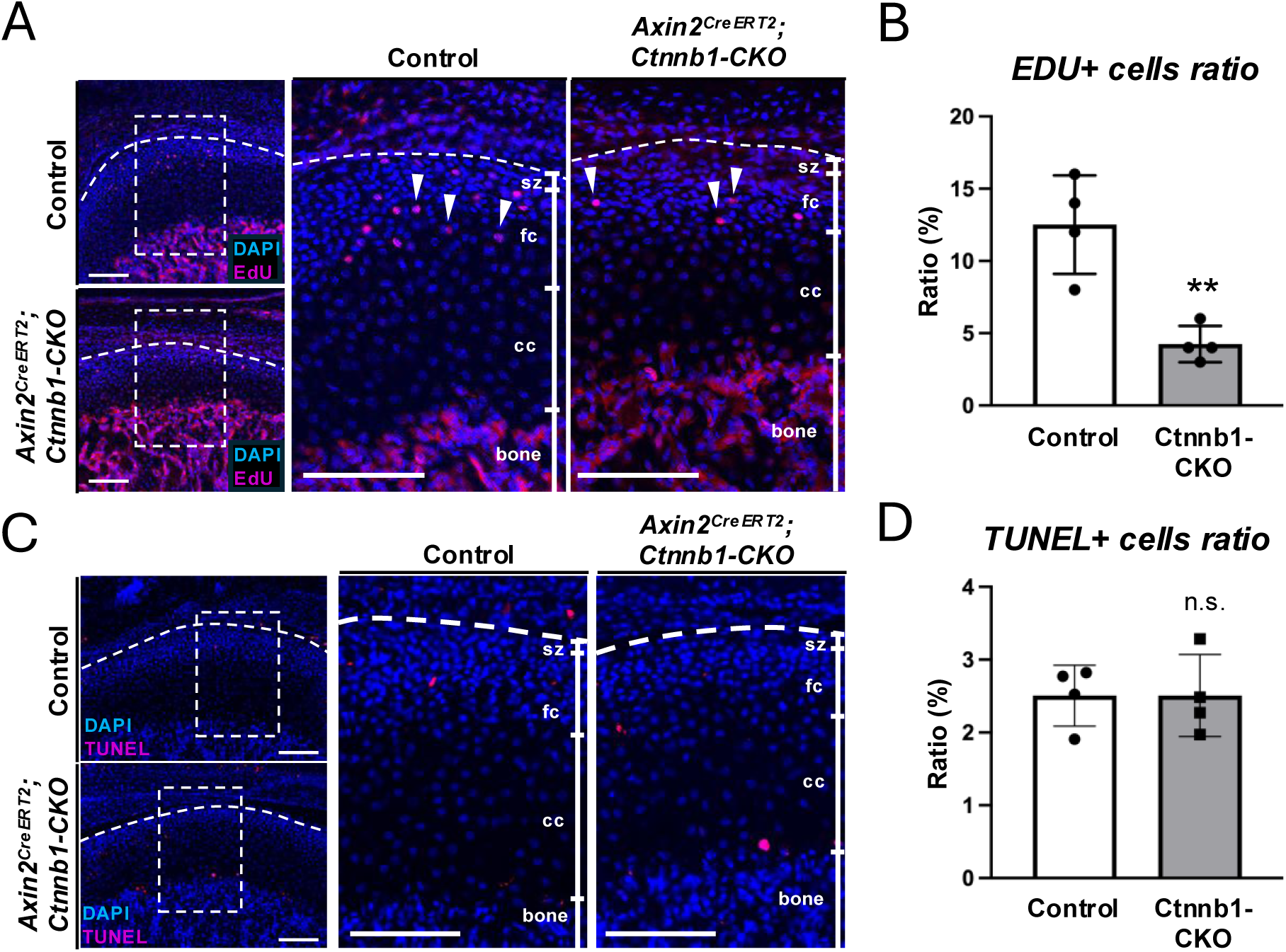
Proliferation and apoptosis analyses following β-catenin deletion. (A) EdU incorporation assay in control and *Axin2^CreERT2^;Ctnnb1^fl/fl^* mice. EdU-positive cells are significantly reduced in mutant fibrocartilage compared with controls, consistent with Ki67 immunostaining results. (B) Quantification of EdU-positive cells within the fibrocartilage compartment. (C) Apoptosis analysis by TUNEL staining. Apoptotic cells are rarely detected in either genotype and show no significant difference between control and mutant condyles. (D) Quantification of TUNEL-positive cells within the fibrocartilage compartment. Data represent mean ± SD from biologically independent mice (n = 4 per genotype). Statistical significance was assessed using two-tailed Student’s t-test. n.s., not significant; **P < 0.01. sz, Superficial zone; fc, fibrocartilage zone; cc, chondrocartilage zone. Scale bars: 100 μm.

**Extended Data Fig. 6.**
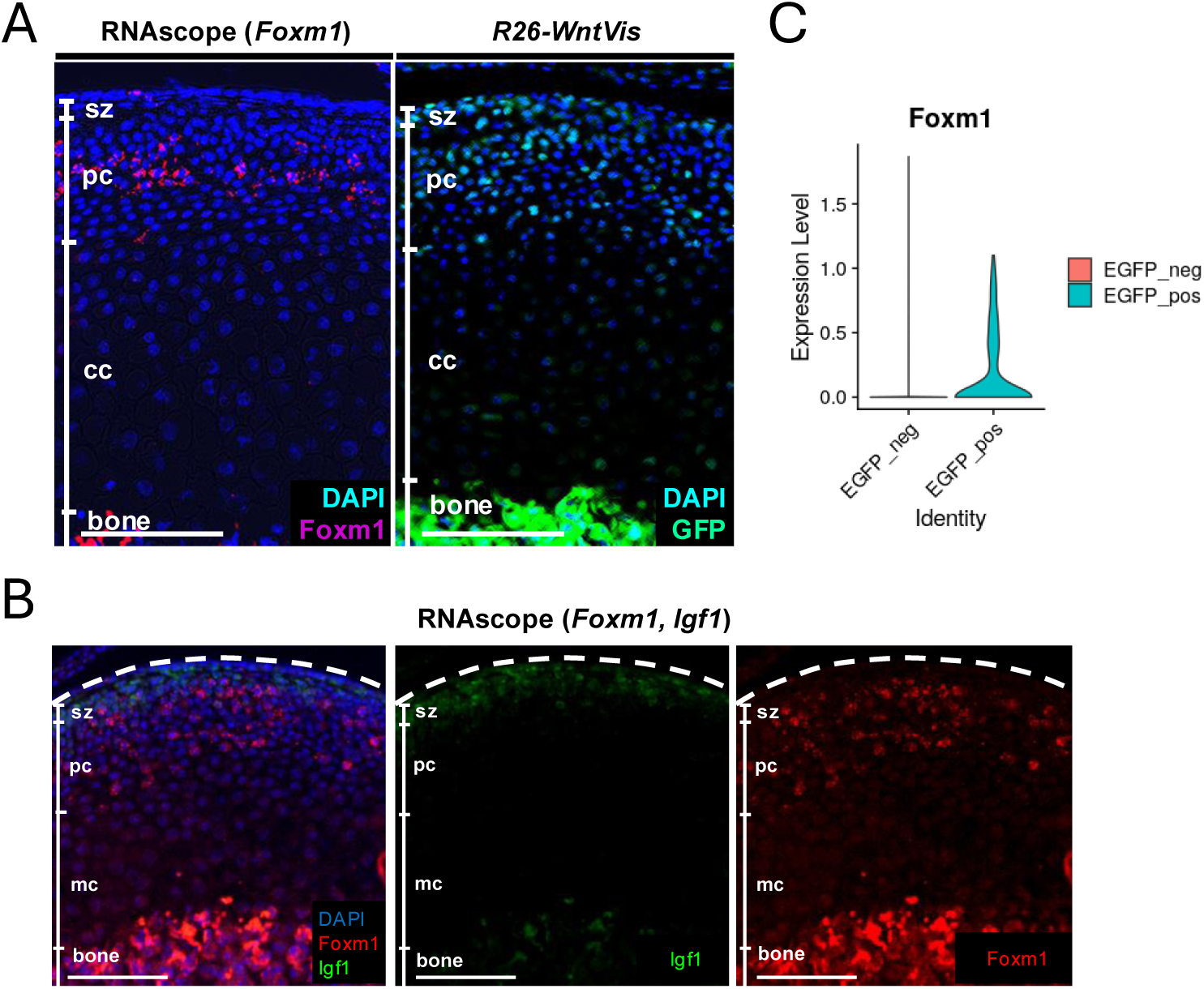
Spatial and quantitative validation of Foxm1 expression in Wnt-responsive fibrocartilage cells. (A) RNAscope in situ hybridization showing *Foxm1* transcript localization within the fibrocartilage compartment of the mandibular condyle. (B) RNAscope detection of *Igf1* transcripts enriched in the superficial region of the fibrocartilage layer. (C) Violin plot comparing *Foxm1* expression between *H2B-EGFP*–positive and *H2B-EGFP*–negative cells within the MC-progenitor cluster. Scale bars: 100 μm.

**Extended Data Fig. 7.**
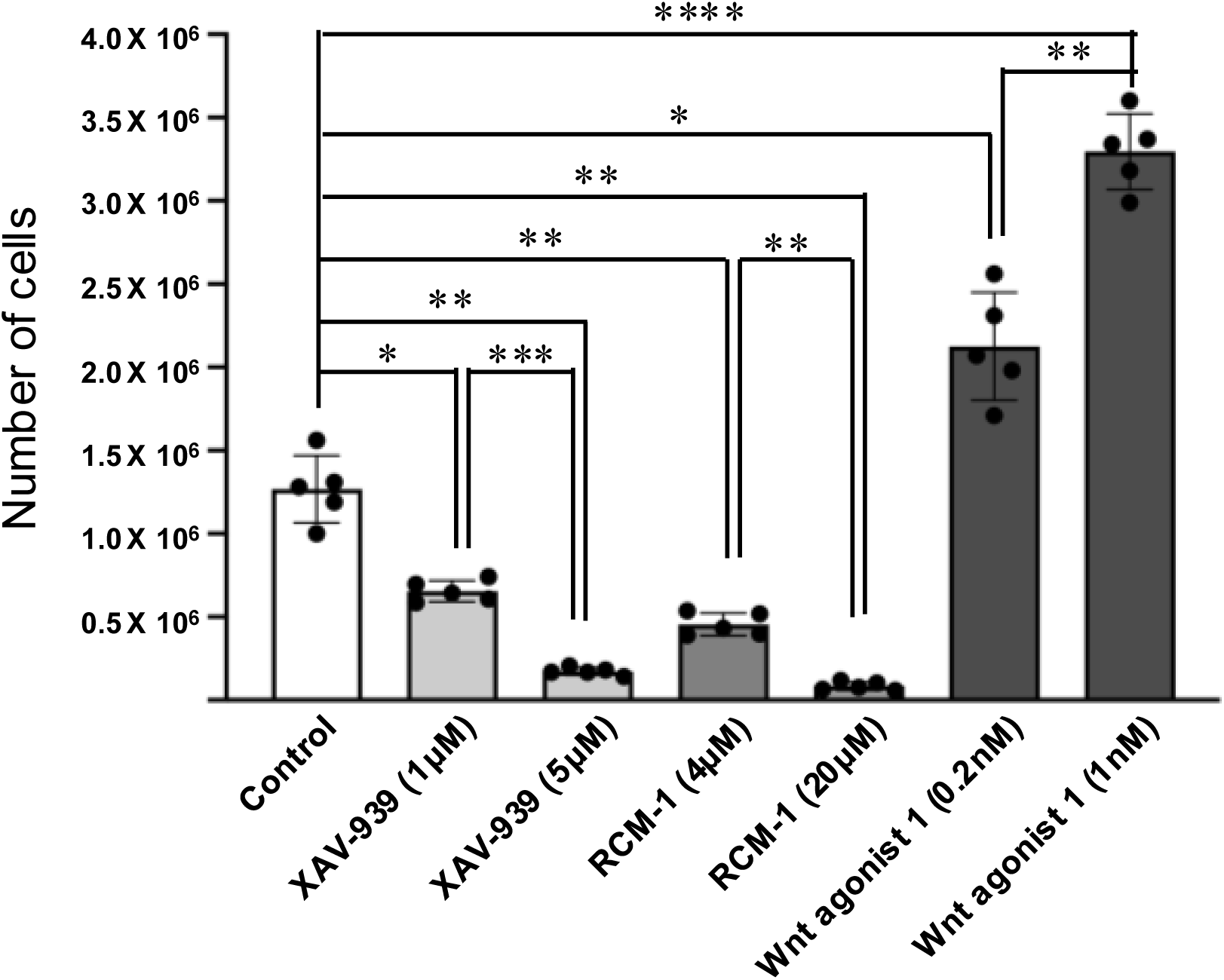
Pharmacological inhibition of Wnt and Foxm1 signaling suppresses proliferation of Wnt-responsive cells. Isolated Wnt-responsive cells were cultured *in vitro* and treated with the Wnt pathway inhibitor XAV-939 (1 μM or 5 μM), the Foxm1 inhibitor RCM-1 (4 μM or 20 μM), or the Wnt agonist Wnt agonist 1 (0.2 nM or 1 nM). Cell proliferation was quantified by measuring total cell number after treatment. Pharmacological inhibition of canonical Wnt signaling using XAV-939 significantly reduced cell expansion compared with control cultures. Similarly, inhibition of Foxm1 using RCM-1 markedly suppressed proliferation. In contrast, activation of canonical Wnt signaling by Wnt agonist 1 increased cell numbers relative to controls. These findings support a cooperative role for canonical Wnt signaling and Foxm1 in promoting proliferative capacity of Wnt-responsive cells *in vitro*. Bars represent mean ± s.d. Each dot represents one biologically independent sample. Statistical significance was determined by one-way ANOVA followed by multiple comparisons testing. *P < 0.05; **P < 0.01; ***P < 0.001; ****P < 0.0001.

**Extended Data Fig. 8.**
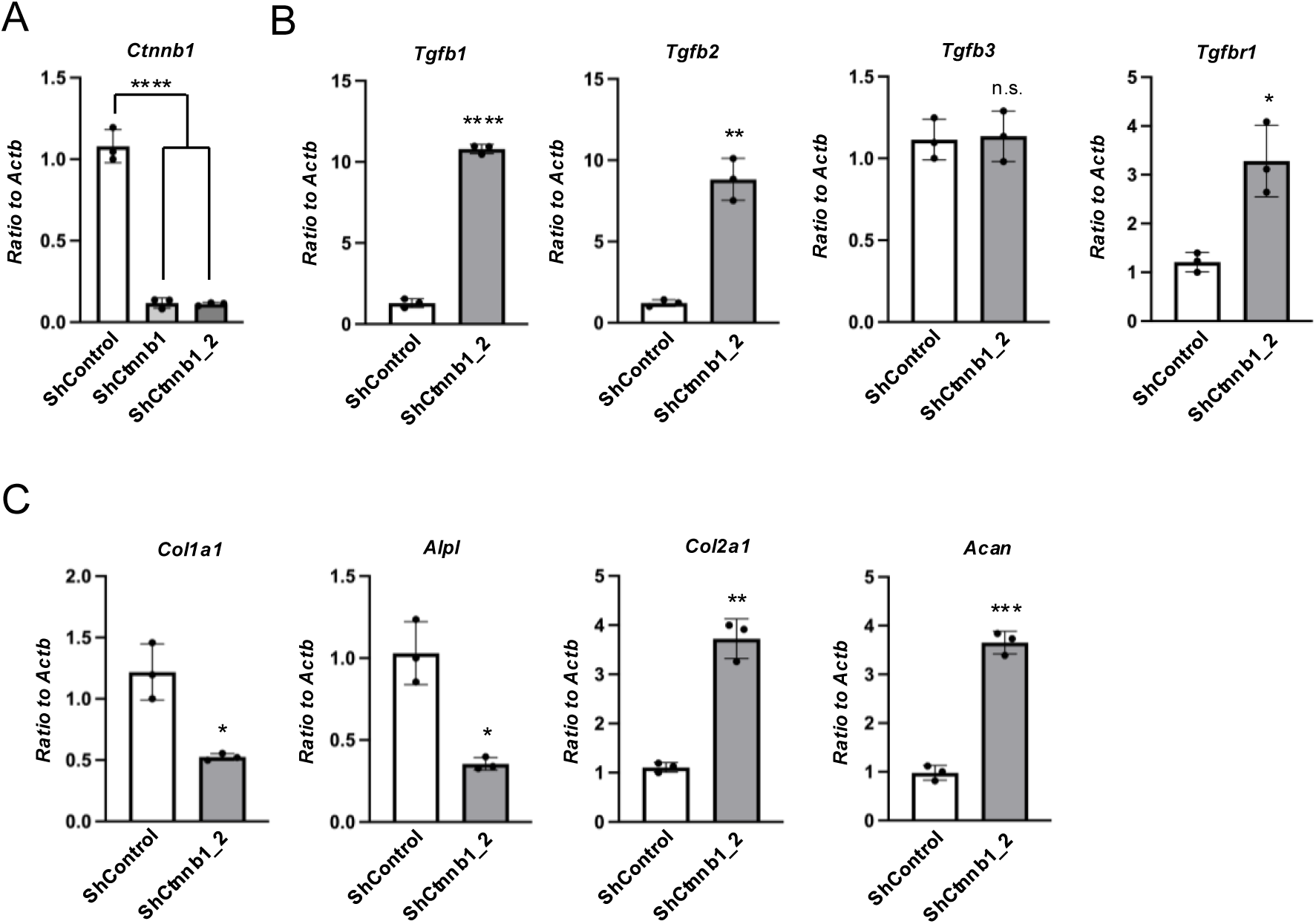
Independent shRNA-mediated knockdown validates Ctnnb1 loss-of-function phenotypes. (A) Quantitative PCR analysis of *Ctnnb1* expression in Wnt-responsive stem/progenitor cells transduced with *control shRNA* (*shControl*) or two independent *shRNAs* targeting *Ctnnb1* (*shCtnnb1 and shCtnnb1_2*). Both *shRNAs* efficiently reduced *Ctnnb1* transcript levels compared with control cells. (B) Quantitative PCR analysis of *Tgfb1*, *Tgfb2*, and *Tgfb3* expression following knockdown of *Ctnnb1* using an independent *shRNA* (*shCtnnb1_2*). Consistent with the primary knockdown experiment shown in Fig. 5, *Ctnnb1* depletion increased *Tgfb1* and *Tgfb2* expression, whereas *Tgfb3* expression remained largely unchanged. (C) Quantitative PCR analysis of osteogenic markers (*Col1a1*, *Alpl*) and chondrogenic markers (*Col2a1*, *Acan*) in cells expressing *shControl* or *shCtnnb1_2*. *Ctnnb1* knockdown enhanced expression of chondrogenic markers while reducing osteogenic markers, consistent with the differentiation phenotypes observed in Fig. 5. Data represent mean ± s.d. from *n* = 3 biologically independent experiments, each analyzed in technical triplicate. Statistical significance was determined by one-way ANOVA followed by multiple comparisons testing (A) and two-tailed Student’s t-test (B, C). n.s., not significant; *P < 0.05; **P < 0.01; ***P < 0.001; ****P < 0.0001.

## Reference

Acri, T. M., Shin, K., Seol, D., Laird, N. Z., Song, I., Geary, S. M., … Salem, A. K. (2019). Tissue Engineering for the Temporomandibular Joint. Adv Healthc Mater, 8(2), e1801236. doi:10.1002/adhm.201801236

Arnett, G. W., Milam, S. B., & Gottesman, L. (1996). Progressive mandibular retrusion--idiopathic condylar resorption. Part I. Am J Orthod Dentofacial Orthop, 110(1), 8–15. doi:10.1016/s0889-5406(96)70081-1

Bergen, V., Lange, M., Peidli, S., Wolf, F. A., & Theis, F. J. (2020). Generalizing RNA velocity to transient cell states through dynamical modeling. Nat Biotechnol, 38(12), 1408–1414. doi:10.1038/s41587-020-0591-3

Bhavanasi, D., & Klein, P. S. (2016). Wnt Signaling in Normal and Malignant Stem Cells. Curr Stem Cell Rep, 2(4), 379–387. doi:10.1007/s40778-016-0068-y

Bryndahl, F., Eriksson, L., Legrell, P. E., & Isberg, A. (2006). Bilateral TMJ disk displacement induces mandibular retrognathia. J Dent Res, 85(12), 1118–1123. doi:10.1177/154405910608501210

Chan, C. K., Seo, E. Y., Chen, J. Y., Lo, D., McArdle, A., Sinha, R., … Longaker, M. T. (2015). Identification and specification of the mouse skeletal stem cell. Cell, 160(1-2), 285–298. doi:10.1016/j.cell.2014.12.002

Chen, Y., Li, Y., Xue, J., Gong, A., Yu, G., Zhou, A., … Huang, S. (2016). Wnt-induced deubiquitination FoxM1 ensures nucleus β-catenin transactivation. EMBO J, 35(6), 668–684. doi:10.15252/embj.201592810

Day, T. F., Guo, X., Garrett-Beal, L., & Yang, Y. (2005). Wnt/beta-catenin signaling in mesenchymal progenitors controls osteoblast and chondrocyte differentiation during vertebrate skeletogenesis. Dev Cell, 8(5), 739–750. doi:10.1016/j.devcel.2005.03.016

Detamore, M. S., & Athanasiou, K. A. (2003). Structure and function of the temporomandibular joint disc: implications for tissue engineering. J Oral Maxillofac Surg, 61(4), 494–506. doi:10.1053/joms.2003.50096

Embree, M. C., Chen, M., Pylawka, S., Kong, D., Iwaoka, G. M., Kalajzic, I., … Mao, J. J. (2016). Exploiting endogenous fibrocartilage stem cells to regenerate cartilage and repair joint injury. Nat Commun, 7, 13073. doi:10.1038/ncomms13073

Enlow, D., & Hans, M. (1996). *Essentials of Facial Growth*. Philadelphia: Saunders.

Gartel, A. L., Ye, X., Goufman, E., Shianov, P., Hay, N., Najmabadi, F., & Tyner, A. L. (2001). Myc represses the p21(WAF1/CIP1) promoter and interacts with Sp1/Sp3. Proc Natl Acad Sci U S A, 98(8), 4510–4515. doi:10.1073/pnas.081074898

Guo, X., & Wang, X. F. (2009). Signaling cross-talk between TGF-beta/BMP and other pathways. Cell Res, 19(1), 71–88. doi:10.1038/cr.2008.302

Harada, N., Tamai, Y., Ishikawa, T., Sauer, B., Takaku, K., Oshima, M., & Taketo, M. M. (1999). Intestinal polyposis in mice with a dominant stable mutation of the beta-catenin gene. EMBO J, 18(21), 5931–5942. doi:10.1093/emboj/18.21.5931

He, T. C., Sparks, A. B., Rago, C., Hermeking, H., Zawel, L., da Costa, L. T., … Kinzler, K. W. (1998). Identification of c-MYC as a target of the APC pathway. Science, 281(5382), 1509–1512. doi:10.1126/science.281.5382.1509

Hill, T. P., Später, D., Taketo, M. M., Birchmeier, W., & Hartmann, C. (2005). Canonical Wnt/beta-catenin signaling prevents osteoblasts from differentiating into chondrocytes. Dev Cell, 8(5), 727–738. doi:10.1016/j.devcel.2005.02.013

Holland, J. D., Klaus, A., Garratt, A. N., & Birchmeier, W. (2013). Wnt signaling in stem and cancer stem cells. Curr Opin Cell Biol, 25(2), 254–264. doi:10.1016/j.ceb.2013.01.004

Inoki, K., Ouyang, H., Zhu, T., Lindvall, C., Wang, Y., Zhang, X., … Guan, K. L. (2006). TSC2 integrates Wnt and energy signals via a coordinated phosphorylation by AMPK and GSK3 to regulate cell growth. Cell, 126(5), 955–968. doi:10.1016/j.cell.2006.06.055

Inubushi, T., Kawazoe, A., Miyauchi, M., Kudo, Y., Ao, M., Ishikado, A., … Takata, T. (2012). Molecular mechanisms of the inhibitory effects of bovine lactoferrin on lipopolysaccharide-mediated osteoclastogenesis. J Biol Chem, 287(28), 23527–23536. doi:10.1074/jbc.M111.324673

Inubushi, T., Lemire, I., Irie, F., & Yamaguchi, Y. (2018). Palovarotene Inhibits Osteochondroma Formation in a Mouse Model of Multiple Hereditary Exostoses. J Bone Miner Res, 33(4), 658–666. doi:10.1002/jbmr.3341

Inubushi, T., Nozawa, S., Matsumoto, K., Irie, F., & Yamaguchi, Y. (2017). Aberrant perichondrial BMP signaling mediates multiple osteochondromagenesis in mice. JCI Insight, 2(15), e90049. doi:10.1172/jci.insight.90049

Kalajzic, Z., Li, H., Wang, L. P., Jiang, X., Lamothe, K., Adams, D. J., … Kalajzic, I. (2008). Use of an alpha-smooth muscle actin GFP reporter to identify an osteoprogenitor population. Bone, 43(3), 501–510. doi:10.1016/j.bone.2008.04.023

Lowry, W. E., Blanpain, C., Nowak, J. A., Guasch, G., Lewis, L., & Fuchs, E. (2005). Defining the impact of beta-catenin/Tcf transactivation on epithelial stem cells. Genes Dev, 19(13), 1596–1611. doi:10.1101/gad.1324905

McNamara, J. A. (1980). Functional determinants of craniofacial size and shape. Eur J Orthod, 2(3), 131–159. doi:10.1093/ejo/2.3.131

Mizuhashi, K., Ono, W., Matsushita, Y., Sakagami, N., Takahashi, A., Saunders, T. L., … Ono, N. (2018). Resting zone of the growth plate houses a unique class of skeletal stem cells. Nature, 563(7730), 254–258. doi:10.1038/s41586-018-0662-5

Nusse, R., & Clevers, H. (2017). Wnt/β-Catenin Signaling, Disease, and Emerging Therapeutic Modalities. Cell, 169(6), 985–999. doi:10.1016/j.cell.2017.05.016

Park, D., Spencer, J. A., Koh, B. I., Kobayashi, T., Fujisaki, J., Clemens, T. L., … Scadden, D. T. (2012). Endogenous bone marrow MSCs are dynamic, fate-restricted participants in bone maintenance and regeneration. Cell Stem Cell, 10(3), 259–272. doi:10.1016/j.stem.2012.02.003

Robinson, J., O’Brien, A., Chen, J., & Wadhwa, S. (2015). Progenitor Cells of the Mandibular Condylar Cartilage. Curr Mol Biol Rep, 1(3), 110–114. doi:10.1007/s40610-015-0019-x

Ruscitto, A., Chen, P., Tosa, I., Wang, Z., Zhou, G., Safina, I., … Embree, M. C. (2023). Lgr5-expressing secretory cells form a Wnt inhibitory niche in cartilage critical for chondrocyte identity. Cell Stem Cell, 30(9), 1179–1198.e1177. doi:10.1016/j.stem.2023.08.004

Schellhas, K. P., Pollei, S. R., & Wilkes, C. H. (1993). Pediatric internal derangements of the temporomandibular joint: effect on facial development. Am J Orthod Dentofacial Orthop, 104(1), 51–59. doi:10.1016/0889-5406(93)70027-L

Shibukawa, Y., Young, B., Wu, C., Yamada, S., Long, F., Pacifici, M., & Koyama, E. (2007). Temporomandibular joint formation and condyle growth require Indian hedgehog signaling. Dev Dyn, 236(2), 426–434. doi:10.1002/dvdy.21036

Takemoto, T., Abe, T., Kiyonari, H., Nakao, K., Furuta, Y., Suzuki, H., … Kondoh, H. (2016). R26-WntVis reporter mice showing graded response to Wnt signal levels. Genes Cells, 21(6), 661–669. doi:10.1111/gtc.12364

Tanaka, E., Detamore, M. S., & Mercuri, L. G. (2008). Degenerative disorders of the temporomandibular joint: etiology, diagnosis, and treatment. J Dent Res, 87(4), 296–307. doi:10.1177/154405910808700406

Tetsu, O., & McCormick, F. (1999). Beta-catenin regulates expression of cyclin D1 in colon carcinoma cells. Nature, 398(6726), 422–426. doi:10.1038/18884

Tuwatnawanit, T., Wessman, W., Belisova, D., Sumbalova Koledova, Z., Tucker, A. S., & Anthwal, N. (2025). FSP1/S100A4-Expressing Stem/Progenitor Cells Are Essential for Temporomandibular Joint Growth and Homeostasis. J Dent Res, 104(5), 551–560. doi:10.1177/00220345251313795

Usami, Y., Gunawardena, A. T., Francois, N. B., Otsuru, S., Takano, H., Hirose, K., … Enomoto-Iwamoto, M. (2019). Possible Contribution of Wnt-Responsive Chondroprogenitors to the Postnatal Murine Growth Plate. J Bone Miner Res, 34(5), 964–974. doi:10.1002/jbmr.3658

van de Wetering, M., Sancho, E., Verweij, C., de Lau, W., Oving, I., Hurlstone, A., … Clevers, H. (2002). The beta-catenin/TCF-4 complex imposes a crypt progenitor phenotype on colorectal cancer cells. Cell, 111(2), 241–250. doi:10.1016/s0092-8674(02)01014-0

Wang, X., Kiyokawa, H., Dennewitz, M. B., & Costa, R. H. (2002). The Forkhead Box m1b transcription factor is essential for hepatocyte DNA replication and mitosis during mouse liver regeneration. Proc Natl Acad Sci U S A, 99(26), 16881–16886. doi:10.1073/pnas.252570299

Zhang, N., Wei, P., Gong, A., Chiu, W. T., Lee, H. T., Colman, H., … Huang, S. (2011). FoxM1 promotes β-catenin nuclear localization and controls Wnt target-gene expression and glioma tumorigenesis. Cancer Cell, 20(4), 427–442. doi:10.1016/j.ccr.2011.08.016

